# Regulatory variation controlling architectural pleiotropy in maize

**DOI:** 10.1101/2023.08.19.553731

**Authors:** Edoardo Bertolini, Brian R. Rice, Max Braud, Jiani Yang, Sarah Hake, Josh Strable, Alexander E. Lipka, Andrea L. Eveland

## Abstract

An early event in plant organogenesis is establishment of a boundary between the meristem and differentiating lateral organ. In maize (*Zea mays*), evidence suggests a common gene network functions at boundaries of distinct organs and contributes to pleiotropy between leaf angle and tassel branch number, two agronomic traits. To uncover regulatory variation at the nexus of these two traits, we used regulatory network topologies derived from specific developmental contexts to guide multivariate genome-wide association analyses. In addition to defining network plasticity around core pleiotropic loci, we identified new transcription factors that contribute to phenotypic variation in canopy architecture, and structural variation that contributes to *cis*-regulatory control of pleiotropy between tassel branching and leaf angle across maize diversity. Results demonstrate the power of informing statistical genetics with context-specific developmental networks to pinpoint pleiotropic loci and their *cis*-regulatory components, which can be used to fine-tune plant architecture for crop improvement.

---

Plant architecture has been an important target of selection in crop domestication and improvement ^1^. The domestication of maize from its wild progenitor teosinte involved major architectural shifts in apical dominance and ear morphology that were pivotal in generating the backbone of a highly productive crop ^2^. Improvements to maize architecture through breeding over the last century have substantially contributed to exponential yield gains; more compact plants with upright leaves and smaller, fewer, or upright tassel branches enable increased planting densities while enhancing photosynthetic efficiency in the lower canopy ^3–6^. These architectural traits are outputs of endogenous developmental programs, intricately connected through gene networks that we are just beginning to unravel.

Variation in plant architecture typically involves changes in placement, number, or orientation of lateral organs, which are initiated from populations of pluripotent stem cells called meristems. Often, this is regulated by meristem determinacy programs ^7^ and the concept of signaling centers acting in boundaries adjacent to meristems was proposed to modulate meristem determinacy and architectural diversity ^8^. During organogenesis, a boundary domain is established between the meristem and differentiating lateral organs to restrict meristem maintenance and organ identity genes to their respective zones ^9^. Leaf angle (LA) in grasses is largely determined by patterning and growth of the ligule and auricles, structures that characteristically define the blade-sheath boundary and together act as a hinge to allow the leaf to recline and absorb sunlight ^9^.

Several genes expressed at initiating ligules are also expressed at the boundaries of other lateral organs, such as developing leaf primordia and tassel branches ^10,11^. Among these are *liguleless* (*lg*)*1* and *2*, which encode SQUAMOSA BINDING PROTEIN (SBP) and bZIP transcription factor (TF)s, respectively. Loss-of-function mutants in these genes have compromised ligule and auricle development, resulting in more upright leaves ^12,13^, but also defects in tassel branch number (TBN) and angle (TBA) (Supplemental Note). Mutants in *lg2* make few to no tassel branches that are upright compared to those of normal siblings ^14^, and *lg1* has significantly fewer branches but the phenotype is less severe than in *lg2* ^15^. Several other maize mutants also show pleiotropic defects between leaf and tassel architecture traits, including those with altered brassinosteroid (BR) signaling, a plant growth hormone that modulates cell division and expansion ^16,17^ (Supplemental Note).

Core gene regulatory modules appear to underlie formation of a boundary, whether it is the boundary at the ligule, the boundary between leaf primordium and meristem, or between tassel branch and rachis. Similar to seminal findings in animal systems ^18^, these common modules have likely been co-opted for development of distinct tissues and underlie pleiotropy found between LA and TBN, important agronomic traits in maize improvement. Although genome-wide association studies (GWAS) identified significant SNP-trait associations for TBN in proximity to *lg1* and *lg2*, pleiotropy between these traits is less prominent in natural populations ^19^. This is likely due to regulatory variation within natural diversity, e.g., *cis*-regulatory elements that specify spatiotemporal patterning of gene expression and are hypothesized to be key drivers of phenotypic variation ^20,21^.

Pleiotropy, the effect of a gene on multiple phenotypic characters, is a major cause of evolutionary constraint, and regulatory variation in pleiotropic loci underpins adaptive evolution and developmental plasticity. Looking forward, the success of new technologies that allow precise engineering of genomes and pathways will depend on our understanding of pleiotropy in gene networks and devising ways of dissociating pleiotropic effects during crop improvement ^22,23^. The pleiotropy that exists between LA and TBN in maize, and the mutants that provide a genetic framework for linking these traits, make it an ideal system for dissecting control points in context-specific gene regulation. Here, we leverage this system and a novel approach that integrates developmental biology, network graph theory and quantitative genetics to identify new factors and regulatory variation that contribute to pleiotropy in tassel and leaf architecture.

## Results

### A transcriptional framework for molecular explorations of tassel branching and leaf angle

To delineate the gene networks underlying tassel branching and ligule development, and tissue-specific rewiring around pleiotropic loci, we leveraged a panel of maize mutants with developmental defects in TBN, LA or both, all well-introgressed into the B73 genetic background (Fig. 1a). Mutants with altered TBN and/or TBA are described in the Supplemental Note and include: *lg1*, *lg2*, *wavy auricle on blade1* (*wab1*) and the dominant *Wab1-R* allele, *ramosa1* (*ra1*), *ra2*, *brassinosteroid insensitive1* (*bri1*)*-RNAi* and *bin2-RNAi*. These genetic stocks, including B73 control plants, were grown in environmentally-controlled chambers to enable precise developmental staging of different genotypes. Immature tassel primordia were hand-dissected immediately before and after primary branch initiation (stage 1 and 2, respectively; Supplementary Fig. 1). We also sampled across a five-stage developmental trajectory from normal B73 tassels; the additional 3 stages representing different meristem transitions after primary branch initiation. Mutants with defects in LA included *lg1*, *lg2*, *Wab1-R*, *bri1-RNAi*, *bin2-RNAi*, and *feminized upright narrow* (*fun*), which makes a ligule but no auricles (Supplemental Note). We sampled two sections of vegetative shoot from these mutants and B73 controls, which we refer to as shoot apex 1 and 2: shoot apex 1 includes the shoot apical meristem (SAM) and cells that are pre-patterned to be ligule, and shoot apex 2 was taken above the meristem to include leaf primordia with developed ligules (Supplementary Fig. 1).

**Fig. 1:**
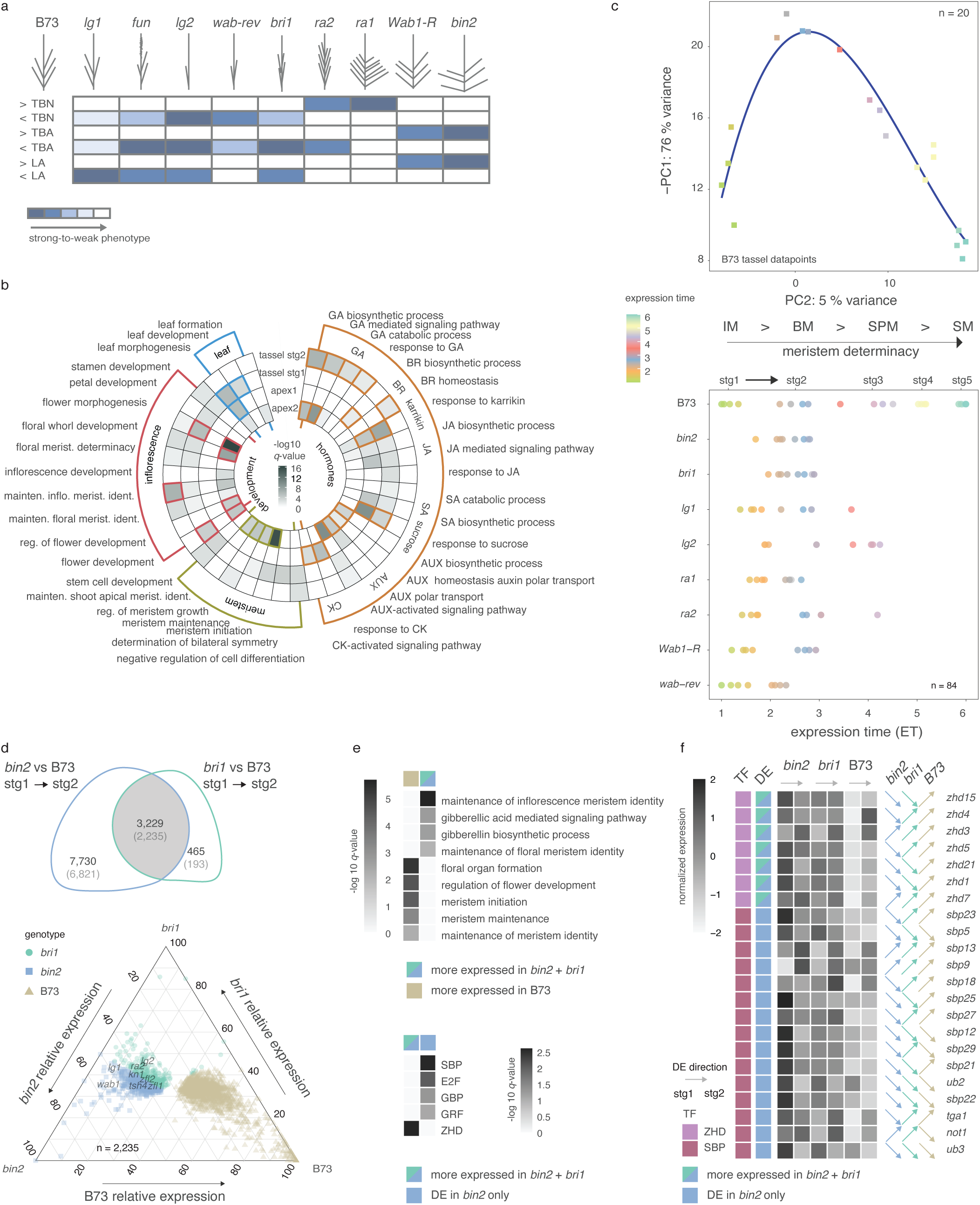
Transcriptional analyses across maize mutants with defects in leaf angle and tassel branching. **a)** Panel of maize mutants used in this study depicting unique tassel branching phenotypes compared to B73 controls. Color scale in the heatmap indicates severity of tassel branch number (TBN), tassel branch angle (TBA) and leaf angle (LA) phenotype deviations from B73 controls; dark blue-to-white = strong-to-weak phenotype. **b)** Enrichment of GO terms associated with DE gene sets identified during the shift from stage (stg) 1 to stg 2 tassel primordia and from shoot apex 1 to shoot apex 2. **c)** A 2-dimensional section of the first and second PC from the normal B73 tassel developmental gradient (n = 20). The blue line represents the b-spline fit with three degrees of freedom modeled within the dataset and used to classify samples relative to their developmental progression coded as expression time (ET). Colored squares represent the 20 tassel samples in ET. The dot plot below represents tassel primordia stgs 1 and 2 from the different mutant backgrounds. Mutant data points are color-coded relative to B73 ET. Stages of meristem development along normal tassel developmental time are indicated: inflorescence meristem (IM), branch meristem (BM), spikelet pair meristem (SPM), spikelet meristem (SM). **d)** Venn diagram shows genes that are DE in tassel primordia during shift from stg 1 to 2 and DE in *bri1-RNAi* and *bin2-RNAi* mutants compared to B73 controls. Those with stronger deviations at stg1 are shown in parentheses. The ternary plot below shows the relative expression of genes commonly mis-expressed between the two mutants compared to B73 at stg 1. Each dot represents a gene and its coordinates indicate relative expression in the three genotypes. Some examples of classical maize genes are noted. **e)** The top heatmap shows GO terms enriched among DE gene sets commonly expressed either higher or lower in the BR mutants compared to B73 controls in stg 1 tassels (from the ternary plot in d); the bottom heatmap shows overrepresented TF classes among genes expressed higher in *bri1* and *bin2* mutants at stg 1 and those expressed higher in *bin2* only compared to B73 controls. **f)** Relative gene expression trends are shown for ZHD and SBP TF family members that were expressed higher in *bin2* and/or *bri1* mutants compared to B73 controls at stg 1. Arrows indicate direction of expression change between stg 1 and 2 tassel primordia.

RNA-seq was used to profile gene expression across 140 samples (Supplementary Table 1). Principal component analysis (PCA) of the normal B73 samples showed clear separation by tissue type and mutant samples cleanly separated by genotype and tissue, indicating distinct transcriptome profiles (Supplementary Fig. 2). Samples across the five stages of B73 tassel development plotted in PCA space revealed a continuous transcriptional gradient reflecting progression through specific axillary meristem types and progression from indeterminate to determinate states. We first examined early shifts in gene expression during primary branch initiation and leaf differentiation in normal B73 plants, which included genes related to meristem identity and determinacy, organ specification, and growth hormone activity (Fig. 1b and Supplementary Fig. 3). We tested for enrichment of functional categories using Gene Ontologies (GO) within differentially expressed (DE) gene sets that were either up-or down-regulated during the shift from meristem to organ differentiation, i.e., between stage 1 and 2 tassels (1,719) and shoot apex 1 and 2 (1,858) (Fig. 1b, Supplementary Tables 2 and 3). As expected, enrichment of several GO categories shifted in common during tassel branch and ligule development, e.g., those related to meristem activity and determinacy and inflorescence morphogenesis, consistent with common sets of genes recruited for boundary establishment and organ development. GO terms related to leaf morphogenesis showed opposite expression between tissues, consistent with the leaf development program being suppressed during tassel branch outgrowth (Fig. 1b). Several plant hormone pathways were overrepresented and tended to show different trends during tassel branch and ligule development. For example, auxin and BR genes showed common expression trajectories in the two developmental programs, whereas jasmonic acid (JA), salicylic acid (SA) and particularly gibberellic acid (GA), showed opposite trends (Fig. 1b). Members of certain developmental TF families showed specific patterns of overrepresentation in tassel branching and leaf development (Supplementary Table 4 and 5). For example, ZINC FINGER HOMEODOMAIN (ZHD) and GRAS family TFs were overrepresented during ligule development, several of which were annotated with GO terms related to GA signaling (Supplementary Fig. 4).

### Heterochronic shifts in gene expression underlie tassel mutant phenotypes

Along the defined expression trajectory of the B73 tassel, we interpolated the mutant tassel expression data to capture deviations from this trajectory, i.e., shifts in heterochrony. We fit a smooth spline regression model using expression values of the 500 most dynamically expressed genes across normal tassel development and used this model to classify samples relative to their transcriptomes, measured in expression time (ET) units (Fig. 1c). Expression profiles of known marker genes in maize inflorescence development (e.g., meristem identity genes) strongly support the ET classifications (Supplementary Fig. 5). Mutant expression data showed clear shifts from normal tassel branch initiation and development, although phenotypic differences were largely not observed at these stages (Fig. 1c). For example, ET classifications of BR signaling mutants, *bri1*-RNAi and *bin2*-RNAi, were notably shifted compared to controls at stage 1 but not stage 2. One interpretation is that transcriptional events modulated by BR signaling during tassel development occur early at branch initiation and become more similar to controls during branch outgrowth. The *bri1* and *bin2* genes encode positive and negative regulators of BR signaling, respectively, and the RNAi mutants display opposite phenotypes for LA and TBA (Fig. 1a; Supplemental Note). BRs play an important role in maintaining boundary domain identity through control of cell division ^24,25^, which is consistent with strong shifts in BR mutant gene expression as boundaries are developed during primary branch initiation.

Shifts in gene expression between tassel primordia at stages 1 and 2 were compared between normal B73 controls and the two BR signaling mutants. Of 3,229 DE genes that showed a differential expression trajectory in both mutants compared to controls, 2,235 showed a larger difference at stage 1 (Fig. 1d, Supplementary Fig. 6). These latter DE genes were enriched for functions related to both meristem maintenance and identity, consistent with important boundary functions, and GA biosynthesis and signaling, suggesting cross-talk between BR and GA pathways (Fig. 1e). Developmentally dynamic genes that were expressed higher in BR signaling mutants at stage 1 were also significantly enriched for ZHD TF family members (*q* = 2.45^e-03^) (Fig. 1f; Supplementary Tables 6, 7, and 8). Notably, we observed significant enrichment (*q* = 1.87^e-02^) of SBP TFs among genes up-regulated in *bin2-*RNAi mutants only during primary branch initiation when compared to normal plants, several of which have been implicated in grass inflorescence development (Fig. 1f) ^26,27^.

### Gene network plasticity around pleiotropic loci in different developmental contexts

To determine transcriptional hierarchies during tassel branch and ligule development we integrated the expression data into ‘tassel’ and ‘leaf’ gene co-expression networks (GCNs) and gene regulatory networks (GRNs). A common set of 22,000 expressed genes was used to generate the two GCNs by weighted gene co-expression network analysis (WGCNA) ^28^. Genes were grouped based on their similar expression patterns into sixteen and eighteen co-expressed modules in ‘tassel’ and ‘leaf’ networks, respectively, which are indicated by color (Supplementary Fig. 7 and Supplementary Tables 9, 10).

To assess the extent of module conservation between the two GCNs, we conducted a co-expression module preservation analysis based on a permutation method (*n* = 1,000; Methods). We found that 11,221 nodes (52% of the commonly expressed genes) were statistically preserved (𝛼 = 0.05, Fisher’s exact test) and that 69% of the ‘tassel’ modules were conserved with one or more sub-modules in the ‘leaf’ GCN (Fig. 2a and Supplementary Table 11). Two ‘tassel’ GCN modules (“brown” and “blue”) were the most preserved with more than 80% module conservation. The ‘tassel’ “brown” module included overrepresented genes associated with regulation of meristem growth (GO:0010075, *q* = 7.74 e^-05^) and maintenance of shoot apical meristem identity (GO:0010492, *q* = 0.0035) (Supplementary Table 12). GO enrichment analysis was performed on conserved ‘tassel-leaf’ GCN sub-modules (Supplementary Table 13). For example, the ‘tassel’ “brown” GCN module included two conserved sub-modules in the ‘leaf’ GCN; i.e., a ‘leaf’ “green” sub-module (nodes = 176, Fisher’s exact test *P* = 5.83 e^-13^) and a ‘leaf’ “red” sub-module (nodes = 591, Fisher’s exact test *P* = 0) that were enriched for genes involved in GA (GO:0009740, *q*= 5.85 e^-05^) and BR (GO:0009742, *q*= 0.03) mediated signaling pathways, respectively.

**Fig. 2:**
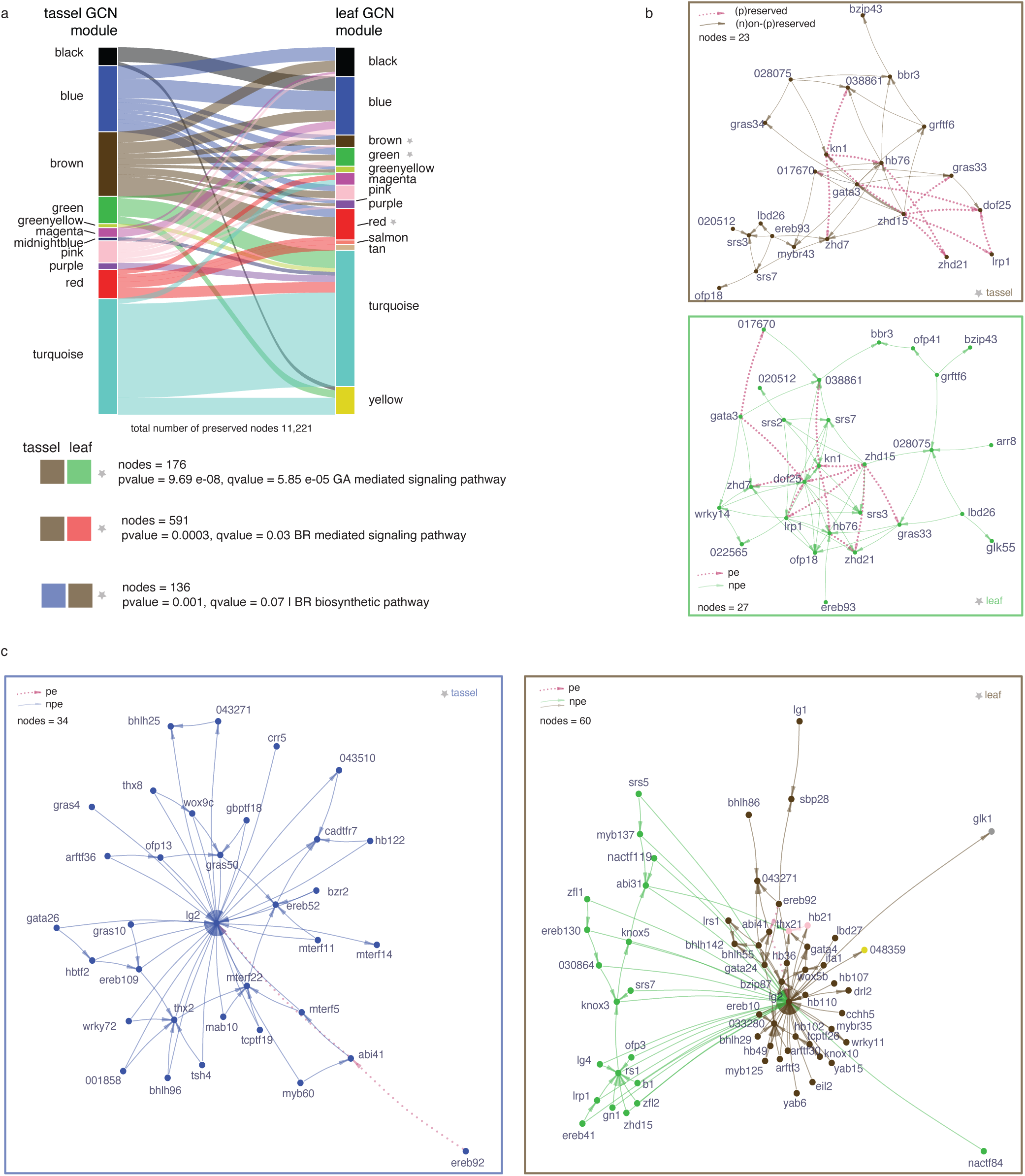
Conserved and divergent network connections between tassel and leaf gene regulatory networks. **a)** Gene co-expression network (GCN) preservation between ‘tassel’ and ‘leaf’ networks. Preserved modules are indicated by color and connected by lines (width is proportional to number of preserved genes). Colored boxes beneath the plot represent example preserved gene co-expression sub-modules. **b)** Topological graph representation of preserved TFs in the ‘tassel’-’leaf’ “brown-green” sub-module based on data from GCN (edge weight) and gene regulatory network (GRN; edge directionality). Pink edges represent preserved regulatory connections and brown or green edges represent the network-specific wires. **c)** Topological graph representation of closest TF neighbors to *liguleless2* (*lg2*) in the ‘tassel’ and ‘leaf’ networks based on data from GCN (edge weight) and GRN (edge directionality). Pink edges represent preserved connections and blue or brown/green edges represent network-specific wires. The edge length is proportional to the weight of gene co-expression. Nodes are colored based on their GCN module designation. Maize gene labels are from MaizeGDB or AGPv4 as Zm00001d_(6)_.

To visualize network rewiring within preserved sub-modules, we conducted a complementary GRN analysis to infer edge directionality. By overlaying the network connections from the GCN and GRN, we observed that several highly-connected regulatory TFs had conserved network connections in ‘tassel’ and ‘leaf’ networks. For example, ZHD15 (Zm00001d003645) showed a high degree of edge conservation in the “brown-green” sub-module suggesting it is a conserved hub gene (Fig. 2b). The TF encoded by *knotted1* (*kn1*), a master regulator of meristem maintenance including regulation of GA pathways ^29^, was also predicted as a conserved hub. In contrast, the uncharacterized TF DOF25 (Zm00001d034163) is potentially a transient hub connected to many genes in the ‘leaf’ GCN (*degree centrality* = 1) but fewer in the ‘tassel’ GCN (*degree centrality* = 0.57).

Since *lg2* mutants are strongly pleiotropic for TBN and LA phenotypes, we investigated tissue-specific connectivity of *lg2* in ‘tassel’ and ‘leaf’ GRNs. We observed substantial rewiring of its closest neighbor nodes with a small degree of edge preservation between ‘tassel’ and ‘leaf’, which included directed edges to *lg2* from TFs EREB92 (Zm00001d000339) and ABI41 (Zm00001d023446) (Fig. 2c). In the ‘tassel’ GRN, *lg2* was co-expressed with and co-regulated by several different TFs including *tassel sheath4* (*tsh4*), an interaction that was validated experimentally ^30^. In the ‘leaf’ GRN, predicted regulators of *lg2* were significantly enriched for HOMEOBOX (HB) (*q* = 1.3e^-04^) and YABBY TFs (*q* = 2.38e^-03^) (Supplementary Table 14), and *lg2* was co-expressed not only with *lg1*, but also *liguleless related sequence1* (*lrs1*) and *sister of liguleless1*, closely related paralogs of *lg2* and *lg1*, respectively. Prior work investigating transcriptional networks at the blade-sheath boundary in maize similarly showed co-expression among these genes ^31^.

### Network motif analysis resolves context-specific topologies and core regulatory factors

To investigate the topology of the GRNs and interconnectedness of TFs, we annotated three-node network motifs; simple, recurrent regulatory circuits that appear in the network at higher frequency than expected by chance ^32^. We used information derived from directed edges inferred in the GRNs and systematically searched for three-node motifs (characterized by at least three edges) that were significantly over-represented in the ‘tassel’ and ‘leaf’ GRNs. Out of ten possible three-node motifs, three types were significantly enriched in both networks (Z score 20) and were present at a higher concentration (≥ 10%), namely the mutual-out, regulating-mutual and feed-forward loop (FFL) (Fig. 3a and Supplementary Table 15). To determine which TFs were predicted to be most influential in the transcriptional circuits, we ranked them based on frequency within annotated three-node motifs. We selected the top one hundred recurrent TFs from each of the ‘tassel’ and ‘leaf’ GCNs (Fig. 3b) for use in subsetting GWAS SNPs (described below). TFs belonging to ZHD (*q*= 0.0028) and HB (*q*= 0.0002) families were the most overrepresented in ‘tassel’ and ‘leaf’ GCNs, respectively.

**Fig. 3:**
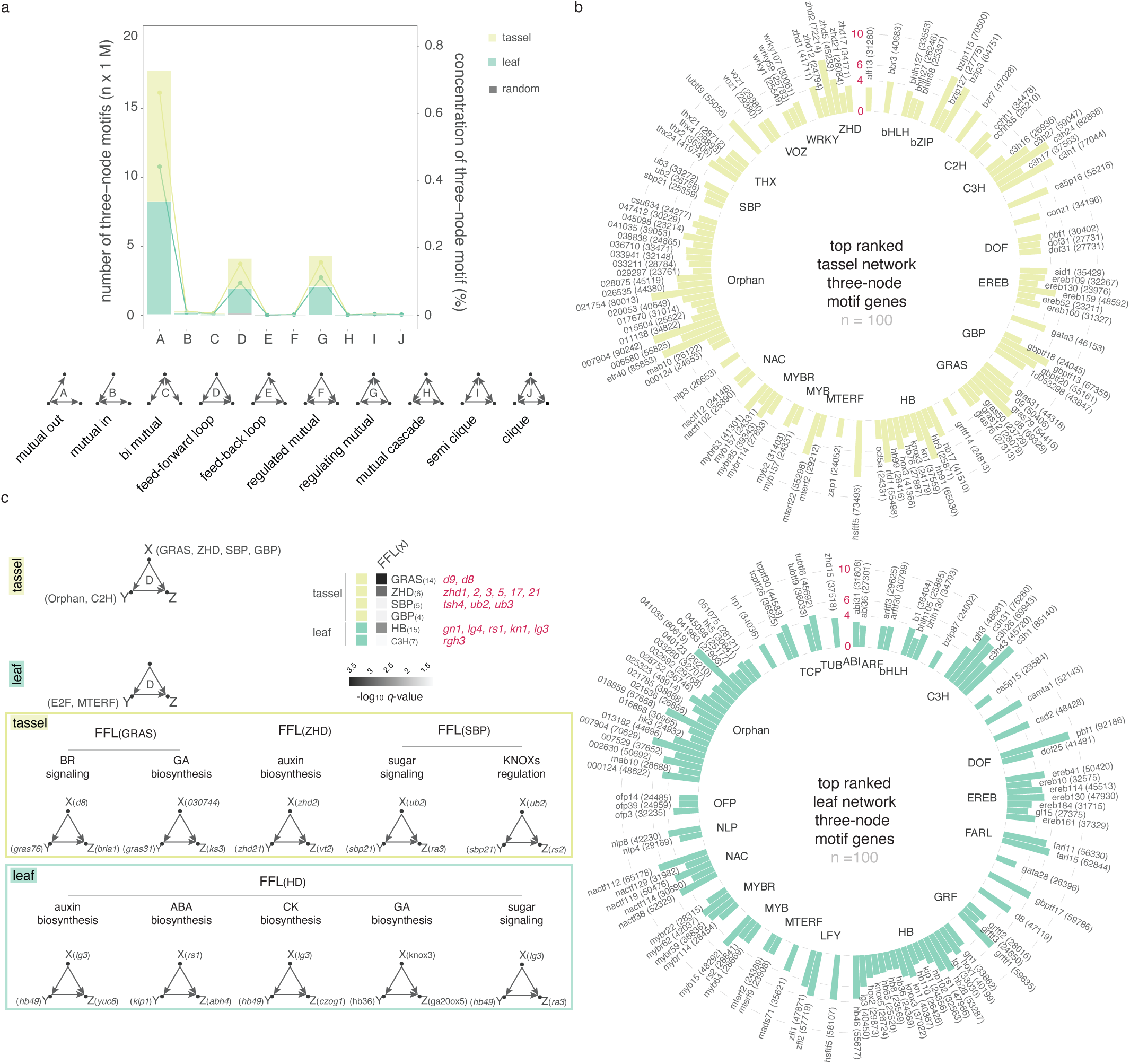
Analysis of three-node network motifs. **a)** The number and type of three-node motifs identified in ‘tassel’ and ‘leaf’ GRNs compared to randomized networks (*n* = 1,000). The x-axis represents the motif type (from A to J) of three-node motifs with edge number ≥ 3. **b)** Top 100 ranked TFs based on their occurrence in the three-node motifs grouped by family. Bars represent TF frequency in the three-node motif (scale in red is TF frequency / 10,000); non-normalized values are within parentheses. TF labels are from MaizeGDB or AGPv4 as Zm00001d_(6)_. **c)** Examples of feed-forward loop (FFL) motifs mediated by select TFs. The heatmap displays overrepresented TFs in FFL_(X)_, (examples of genes in each TF family are noted). Nodes represent genes, while the directed edges represent the potential regulatory relationships.

FFLs have been described across organisms as local, repeated and adapted genetic circuits ^33^. To gain insight into the modularity of the FFLs within our context-specific GCNs, we identified the overrepresented TFs in the two parallel regulation paths controlled by nodes “X” and “Y” (Fig. 3c). GRAS, ZHD and SBP were the most overrepresented TF families in the ‘tassel’ FFL_(X)_ while HBs were overrepresented in the ‘leaf’ FFL_(X)_. In both networks, these TFs were predicted to regulate several genes related to hormone biosynthesis and signaling and nutrient sensing.

### A genomic selection approach to optimize phenotyping of maize diversity for architectural pleiotropy

We took a quantitative approach to identify loci that link TBN and LA in maize. To optimize selection for maximizing diversity in these architecture traits, a genomic best linear unbiased predictions model (GBLUP) was fit using the Goodman-Buckler diversity panel ^34^ and the predicted genomic estimated breeding values (GEBVs) for the following traits in 2,534 diverse lines from the Ames inbred panel ^35^: TBN, LA, ear row number (ERN; included to maximize diversity for ear traits too), and first and second Principal Components of the phenotypes (PhPC1 and PhPC2, respectively) (Supplementary Table 16). Ames lines (*n*=1,064) were randomly selected to represent the distribution of the predicted PhPC1 and manually phenotyped for TBN and LA. Overall, the large portion of heritable variation among the selected genotypes resulted in prediction accuracies of 0.58 and 0.50 for TBN and LA, respectively.

PhPC1 of the selected lines explained 56% of the total variance of TBN and LA, and was explained by trait loadings that were both positive. PhPC2 explained the remaining variance (44%) with positive (LA) and negative (TBN) loadings (Fig. 4a). Interestingly, PhPC1 explained a larger portion of the variance than either TBN or LA, suggesting that PhPC1 may be a better source for detecting pleiotropy than testing phenotypes independently.

**Fig. 4:**
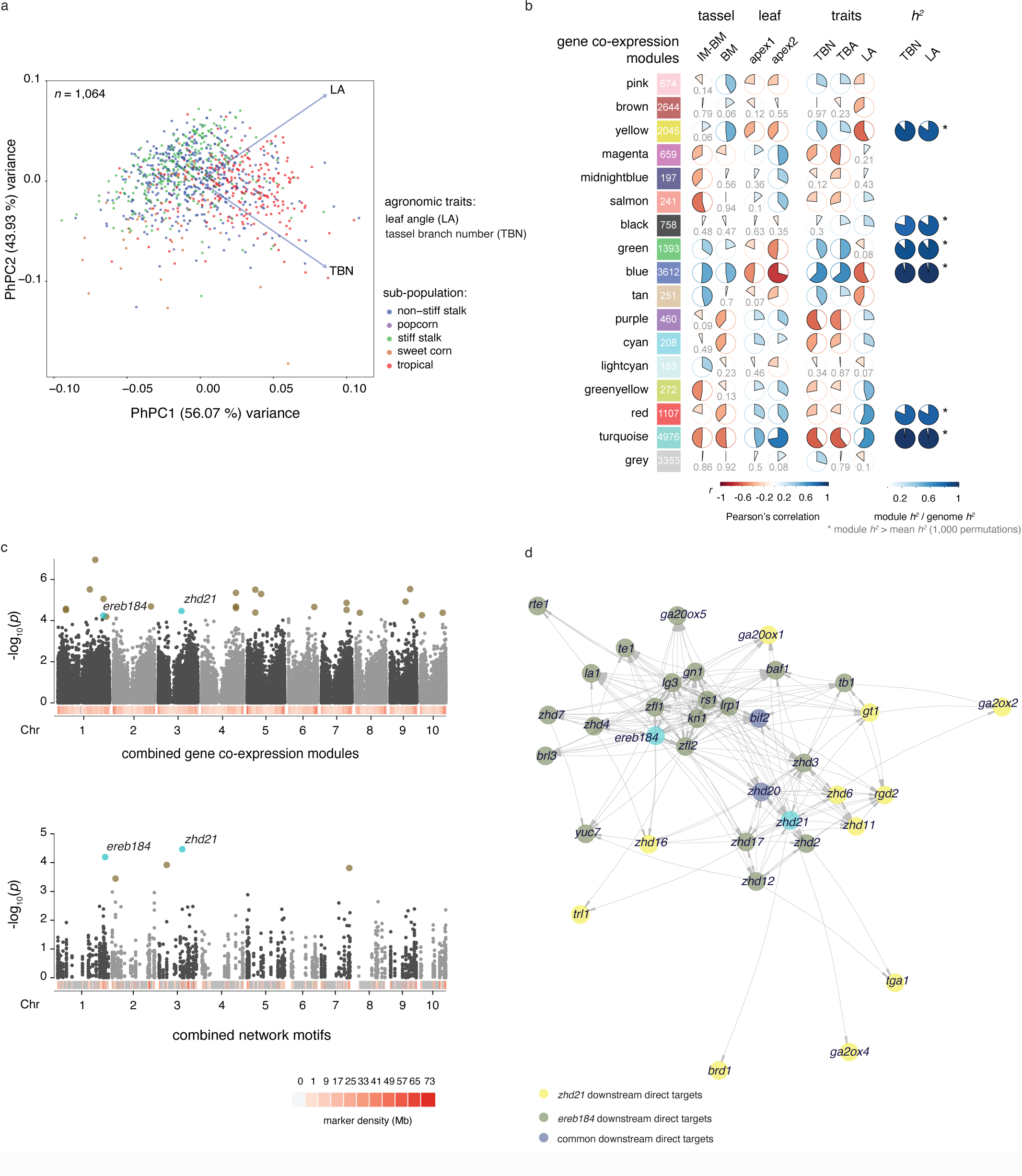
Multi-trait GWAS detected candidate pleiotropic genes for TBN and LA. **a)** Phenotypic Principal Components (PhPCs) of TBN and LA. Individuals are plotted according to their PhPC1 scores of TBN and LA on the x-axis and their PhPC2 scores on the y-axis. The percentage of total variance explained by a component is listed in parentheses on the axis. Blue arrows represent trait loadings for a given PhPC where its direction on the x-axis represents its contribution to PhPC1 and direction on the y-axis, its contribution to PhPC2. Subpopulations are color-coded and grouped according to PhPCs. **b)** Analysis of module-trait relationships for the combined GCN. Modules are represented as colored boxes with the number of co-expressed genes indicated. The correlogram to the right represents the module-trait relationship where shades of blue and red represent positive and negative Pearson’s correlation, respectively, with darker colors indicating a stronger positive or negative correlation. Modules representing more than 80% of the whole genome heritability (*h^2^*) for TBN and LA are indicated. **c)** Multi-trait GWAS results using subsets of markers within 2 kb proximity of genes in the six co-expression modules with significant *h^2^* in panel b (top) and in the top 200 interconnected TFs within three-node motifs from Fig. 3b. Brown and blue dots represent significant SNP-trait associations; blue dots indicate loci identified in both analyses. **d)** Topological graph representation of a sub-network including *ereb184*, *zhd21* and their predicted downstream target genes based on data from the ‘leaf’ GRN (edge directionality).

Because plant architecture has played a key role in maize adaptation, we hypothesized that genes underlying inflorescence and leaf architecture traits would be confounded with population structure. This hypothesis was underscored by how clearly the proxy traits PhPC1 and PhPC2 subdivided the lines into their respective subpopulations (Fig. 4a). Moreover, we observed a moderate correlation (0.53) between the PhPC1 (Supplementary Table 17) and published flowering time data ^36^. These results collectively suggest divergence between TBN, LA, and their pleiotropic regulation across maize subpopulations. Thus, the models we used in the ensuing analysis corrected for both population structure and familial relatedness (Methods).

### Network-assisted multi-trait GWAS identifies genetic loci associated with TBN and LA

SNP markers were prioritized using gene sets related to TBN and LA based on network analyses (described below) and were used to guide multivariate GWAS. While this approach significantly reduced the marker density to varying degrees, we hypothesized that biological information derived from the networks could resolve novel SNP-trait associated markers.

To select genes for marker sub-setting, we generated a comprehensive GCN using the entire RNA-seq dataset (*n*=140) and identified sixteen modules of co-expressed genes. Using a module-feature relationship analysis in WGCNA, we identified modules that had significant correlations with tissue types, developmental stages or agronomic traits (Fig. 4b and Supplementary Table 18). For each GCN module, we estimated TBN and LA narrow-sense heritability (*h^2^*) and compared the observed values to empirical distributions. Modules that fit the following three conditions were selected (*n*= 6): i) significant module-trait correlation, ii) observed *h^2^* values greater than the mean of their respective empirical distribution and iii) at least 80% of genome-wide *h^2^* explained (Fig. 4b and Supplementary Table 19). SNPs overlapping with genes in the selected six co-expression modules (*n*=106,790; within 2 kb upstream and downstream of the gene model) accounted for nearly all of the genome-wide *h^2^* associated with TBN and LA (Supplementary Fig. 8). In addition to the combined GCN, we also used a reductionist approach by selecting the top one hundred recurrent TFs within the three-node network motifs derived from the ‘tassel’ and ‘leaf’ GRNs (Fig. 3b) as an alternative way to partition SNP markers. Remarkably, SNPs (*n*=1,972) overlapping this group of genes still explained a significant portion of the genome-wide *h^2^*, 57% and 70% respectively for TBN and LA (Supplementary Fig. 8).

For each set of markers, from 1) combined GCN and 2) three-node motif analysis, we conducted single-trait GWAS for TBN, LA, and the first two PhPCs, and multi-trait GWAS analyses for TBN and LA. Our GWAS models collectively yielded 71 and 172 non-overlapping single trait-associated SNPs for the combined GCN and three-node motif sets, respectively (Supplementary Table 20). The majority (72%) of the trait-associated markers in the three-node motif set were associated with PhPC2, which represents opposite trait loadings. Genes associated with PhPC2 were significantly enriched with GO terms implicated in meristem determinacy and maintenance (Supplementary Table 21).

Multivariate analysis with the two marker sets identified a total of 23 potential pleiotropic loci where SNPs simultaneously associated with both TBN and LA phenotypes (Fig. 4c and Supplementary Table 22). Two of these SNPs were common in analyses with both marker sets indicating high-confidence associations that may contribute to phenotypic pleiotropy. Both SNPs were located within genes encoding TFs; one in the last exon of *ereb184* (Zm00001d034204), an ortholog of *AINTEGUMENTA1* in *Arabidopsis thaliana* known to control plant growth and floral organogenesis ^37^, and one in the last exon of *zhd21* (Zm00001d041780), which was shown to express in leaf primordia during ligule initiation ^31^. These two markers were also associated with TBN in the single-trait GWAS. Also, the marker in *ereb184* associated with PhPC1 in the single-trait GWAS, reinforcing the notion that PhPC1 may be used to detect pleiotropic loci.

We found that *ereb184* and *zhd21* were connected in the ‘leaf’ GRN through four TFs: *ereb93*, *zhd20, zhd16* and *barren inflorescence2;* the latter regulates tassel branch outgrowth in maize ^38^. Predicted downstream targets of these two TFs suggest that they regulate architecture traits through different but interconnected developmental circuits made of several other *zhd* genes, *knox* genes and genes regulating hormone metabolism, transport and signaling, especially in auxin, GA and BR pathways (Fig. 4d and Supplementary Table 23). The link between *ereb184* and *zhd21* was also supported by statistical epistasis analysis (Supplementary Fig. 9 and Supplementary Table 24), which highlighted their interaction together with other TFs and members of the ZHD family, such as *zhd1, zhd12 and zhd15*.

### Maize network data guide explorations of SNP-trait associations in sorghum

We further tested whether the small set of markers selected based on three-node network motifs in maize was sufficient to guide GWAS to candidate SNP-trait associations for LA in sorghum. Among the maize genes within the motif set, we identified 146 sorghum orthologs and their associated SNPs from the Sorghum Association Panel (SAP) ^39^. Using the selected SNP markers together with publicly available LA phenotype data for the SAP ^40^, we first tested *h^2^*for this trait and found that variation in these genes explained more than 60% of the sorghum genome-wide LA *h^2^* (Fig. 5a). Therefore, we conducted a single-trait GWAS for LA. Eight sorghum LA-associated markers were found within or proximal to seven genes (Fig. 5b and Supplementary Table 25), including orthologs of maize genes *nac112* (Sobic.009G143700), *ofp39* (Sobic.008G042200), *c3h16* (Sobic.006G256500) and Zm00001d026535 (Sobic.006G254500), which were also identified in maize single-trait GWAS for PhPC1, PhPC2 and/or TBN. Additionally, a significant SNP was identified within the maize ortholog of *ereb114* (Sobic.002G022600), a paralog of *ereb184*. Based on first neighbors’ connectivity in the maize ‘leaf’ GRN, *ereb114* is situated either up- or downstream of known *knox* genes involved in meristem maintenance and axillary meristem formation in maize, e.g., *kn1*, *gn1*, *rs1* and *lg4*, suggesting conservation of developmental circuits controlling LA between maize and sorghum (Fig. 5c). Furthermore, *ereb114* is predicted to directly target *zhd21*, another high-confidence gene candidate from the multi-trait GWAS for LA and TBN.

**Fig. 5:**
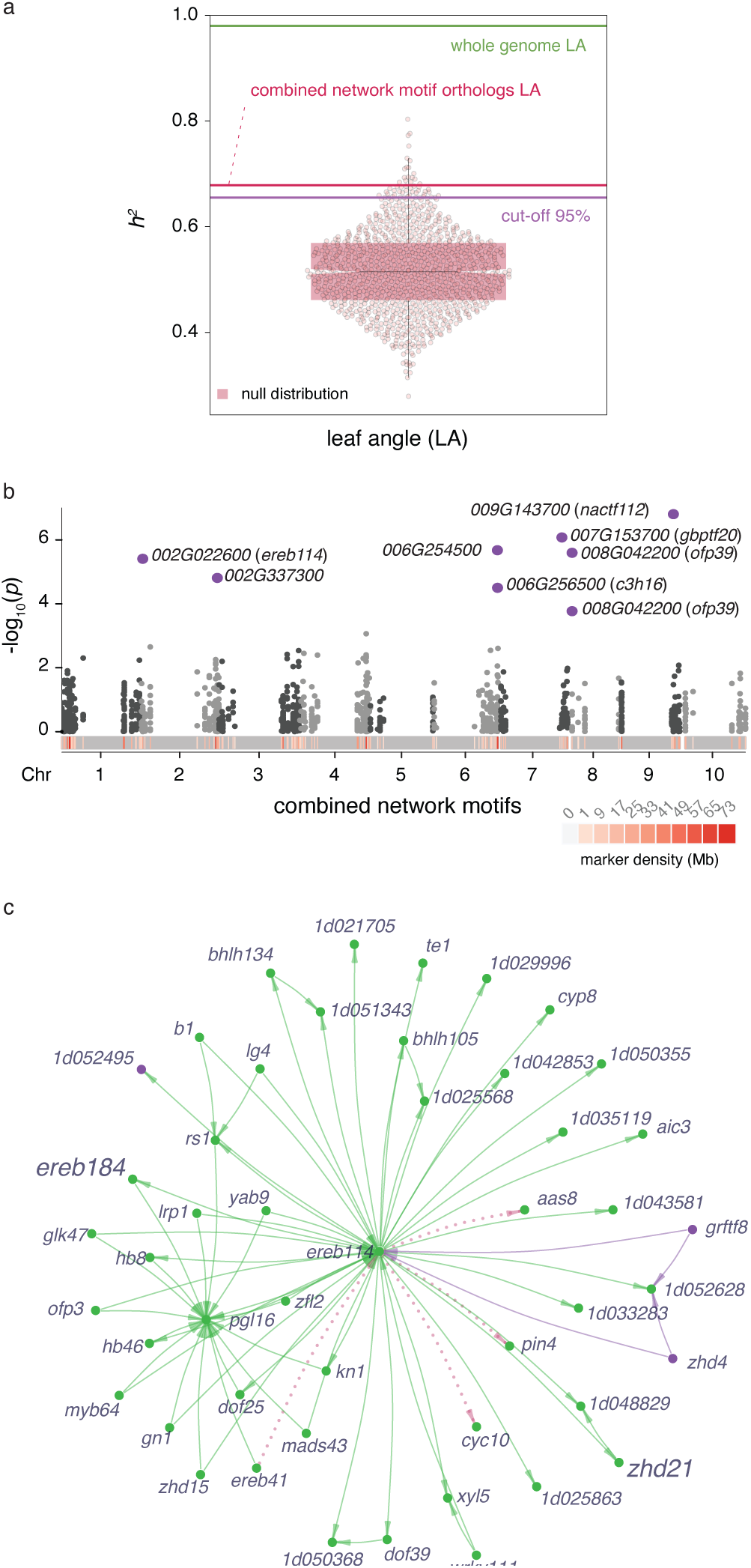
GWAS for LA in sorghum using biological information derived from maize. **a)** LA narrow-sense heritability of 146 sorghum-maize syntenic orthologs in comparison to a null distribution (violin plot). Purple line represents 95% cut-off, red line represents LA *h^2^* of the sorghum orthologs, green line is the whole genome LA *h^2^*. **b)** Manhattan plot showing GWAS results for LA based on markers in proximity of the 146 sorghum-maize syntenic orthologs. Purple dots represent significant SNP-trait associations for LA. **c)** Topological graph representation of *ereb114* and its closest network neighbors in the maize ‘leaf’ GCN (edge weight) and GRN (edge directionality). Green and purple nodes and edges are colored based on module assignment in the ‘leaf’ GCN. Pink edges represent preserved connections with the ‘tassel’ GCN. Edge length is proportional with the gene co-expression weight.

### *zhd* genes modulate tassel and leaf architecture

Throughout this study, genes encoding ZHD TFs repeatedly emerged as important players in tassel branch and ligule development. These TFs appear as putative hubs in the networks or highly connected to known developmental pathways. *Zhd* genes are enriched within BR signaling-responsive genes during early development, and *zhd21* was identified as a high-confidence candidate gene in multi-trait GWAS for TBN and LA. The extensive interconnectedness among numerous ZHD family members suggests functional redundancy and potential for molecular fine-tuning. To test whether disruption of network candidate *zhd* genes have a phenotypic effect on tassel and/or leaf architecture, we studied the effects of independent *Mutator* (*Mu*) transposon insertions in the coding regions of *zhd21* and *zhd1* (UniformMu, Methods). One allele of *zhd21* (*zhd21-1*; mu1056071) showed significantly fewer tassel branches and a more upright LA compared to W22 normal plants (Wilcoxon *p*-value TBN=2.1e^-04^, LA=8.5e^-09^), while a different allele of *zhd21* (*zhd21-2*; mu1018735) and *zhd1* (mu1022277) showed no significant difference in either trait (Fig. 6a). The mutation in *zhd21-1* disrupts the zinc-finger domain, which we expect confers the observable phenotype. Since we hypothesize some degree of functional redundancy among *zhd* genes, lack of phenotypes for the other alleles was not surprising.

**Fig. 6:**
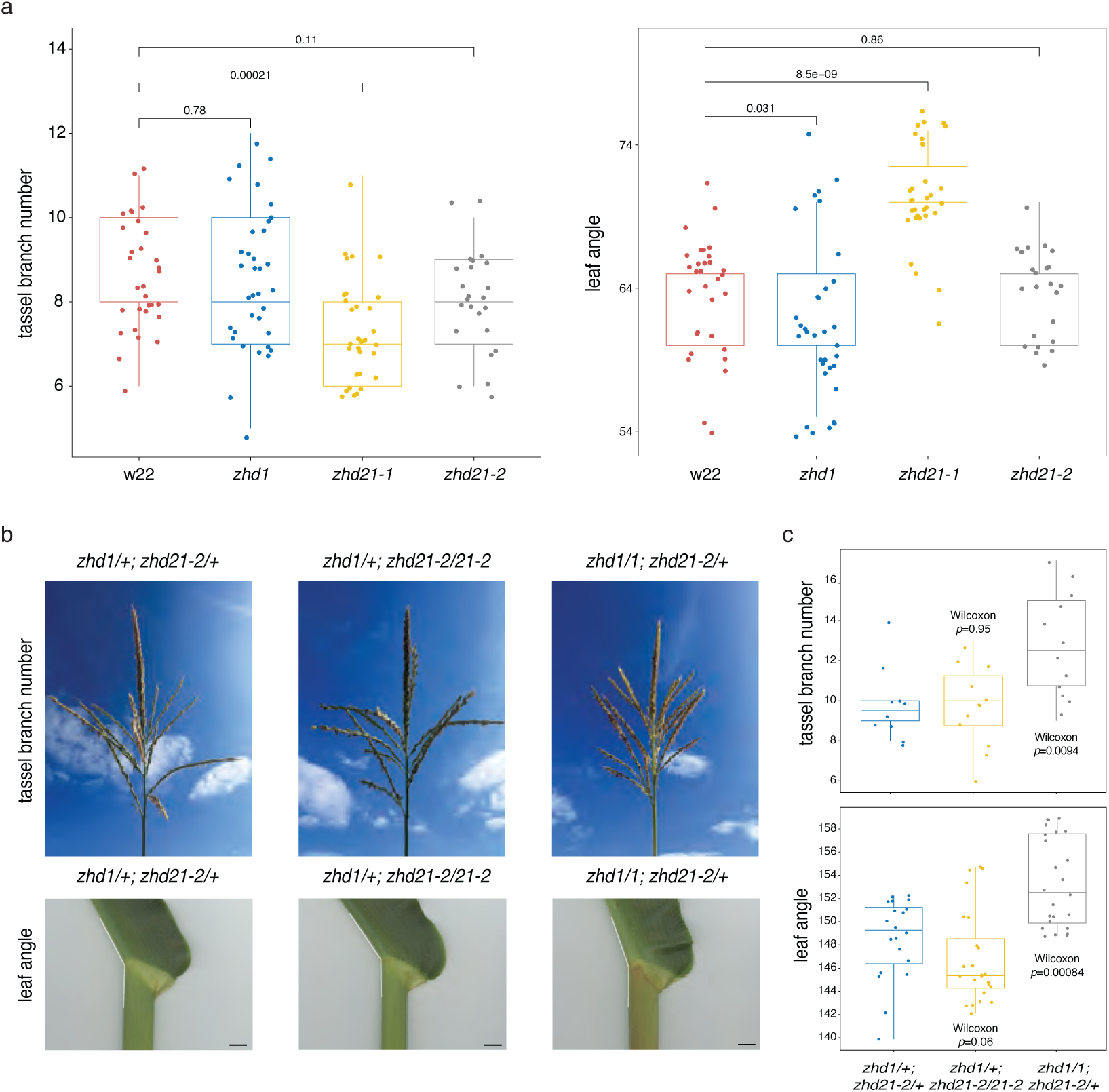
*zhd* mutant alleles contribute to tassel and leaf architecture. **a)** Phenotypic characterization of TBN and LA in homozygous *zhd* alleles compared with w22 control plants. **b**) Representative tassel branch and leaf angle phenotypes resulting from genetic interactions between mutant alleles of *zhd1* and *zhd21*, from left to right: *zhd1/+*;*zhd21-2/+*, *zhd1/+*;*zhd21-2/zhd21-2*, *zhd1/1*;*zhd21-2/+*. Black bar = 1cm. **c)** Among combinations of mutant alleles, *zhd1/1*;*zhd21-2/+* showed significant differences in TBN and LA compared to *zhd1/+*;*zhd21-2/+* normal siblings based on Wilcoxon rank-sum test.

We tested whether stacking *zhd21* and *zhd1* alleles with no apparent phenotypes in homozygous single mutants resulted in architectural differences. Interestingly, plants that were homozygous for the *zhd1* mutant allele and heterozygous for *zhd21-2* had significantly more tassel branches and more upright LA compared to plants heterozygous for both alleles (Wilcoxon *p*-value TBN= 0.0094, LA= 0.0008) (Fig. 6b,c). Plants homozygous for *zhd21-2* and heterozygous for *zhd1* showed no significant difference in either trait. Our results suggest that *zhd21* and *zhd1* influence tassel branching and leaf architecture, and that complex network connectivity among *zhd* family members may allow for precise modulation of pleiotropy in these traits through combinations of alleles.

### Structural variation in the promoter of *ereb184* modulates pleiotropy in TBN and LA

Our gene candidate *ereb184* was among the top hundred recurring TFs within ‘leaf’ GRN three-node motifs and positioned as a regulatory factor in many FFLs (FFL_(x)_; n=7,422). Notably, these FFLs were overrepresented by ZHDs at the FFL_(y)_ node (Fig. 7a). We found that *ereb184*-regulated FFLs potentially regulate a set of downstream genes (FFL_(z)_) associated with functional categories that overlapped with those enriched during tassel branch and ligule development (Fig. 7a and Supplementary Table 26). For example, four KN1-regulated FFLs (validated by ChIP-seq data ^29^) included *ereb184* connections to several known homeotic genes that were predicted in the ‘leaf’ GRN to directly target *lg2* (Fig. 7b).

**Fig. 7:**
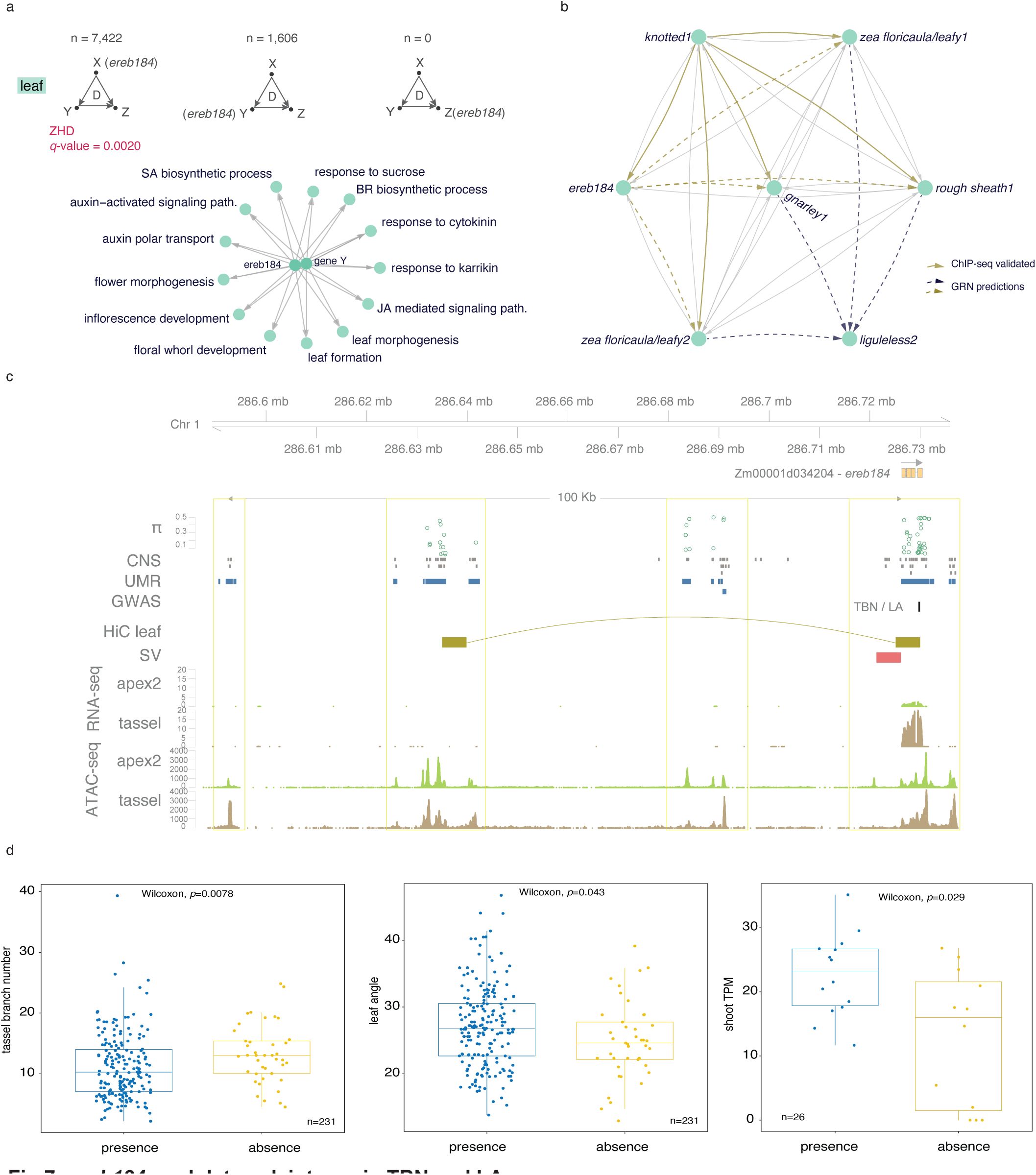
*ereb184* modulates pleiotropy in TBN and LA. **a)** FFLs from the ‘leaf’ GRN are represented with *ereb184* in each of the node positions and number of times each occurs. The graph below shows GO terms overrepresented within the Z position when EREB184 is in the X position. **b)** An example of a sub-network from the ‘leaf’ GRN shows *ereb184* participating in multiple FFLs with KN1 and other proximal-distal patterning TFs that target *lg2*. The solid yellow arrows represent ChIP-seq validated connections. **c)** Genomic view of the intergenic space upstream (100 kb) and just downstream of *ereb184*. The tracks from top to bottom: nucleotide diversity (π), conserved non-coding sequences, multi-trait GWAS, unmethylated regions, leaf chromatin interaction data (Hi-C), structural variation, RNA-seq, and ATAC-seq. Data from public sources are indicated in Data Accessibility. **d)** The effect of structural variation (SV) presence/absence in the promoter region of *ereb184* across 231 accessions on TBN and LA traits, and the effect of SV on *ereb184* expression within 26 NAM founder lines.

Scanning the gene regulatory space around *ereb184,* we observed approximately 100 kb of intergenic sequence upstream of the TSS. We integrated publicly available omics’ datasets to investigate potential regulatory regions (see Data accessibility). We identified three genomic regions with regulatory signatures including unmethylated marks, which colocalized with conserved non-coding sequences and showed increased nucleotide diversity, similar to that overlapping the coding region of *ereb184* (Fig. 7c). To investigate tissue-specific chromatin accessibility, we generated Assay for Transposase-Accessible Chromatin (ATAC)-seq data from the B73 shoot apex 2 tissue (including developed ligules) and compared the profiles in this region to publicly available ATAC-seq data from tassel primordia ^41^. All three regulatory regions showed tissue-specific chromatin signatures that overlapped with other epigenetic marks, highlighting potential tissue-specific regulation of *ereb184*. Overlaying publicly available maize leaf chromatin interaction data (Hi-C) ^42^ predicted chromatin looping between the *ereb184* promoter and one of the upstream intergenic regions (Fig. 7c).

Using public data from the maize Nested Association Mapping (NAM) Consortium ^43^, we identified structural variation (SV) of approximately 5 kb in the promoter region of *ereb184*. Based on resequencing data of the 26 NAM founder lines, this SV was either present or absent, and mainly absent in tropical inbred lines. We also observed tissue-specific chromatin accessibility adjacent to the SV, suggesting it is located within a regulatory region. We tested whether presence/absence variation (PAV) of the SV in the promoter of *ereb184* influenced pleiotropy between TBN and LA. In addition to the NAM founder reference genomes, we extended the analysis to include 231 accessions that we phenotyped from the Goodman-Buckler panel ^34^, incorporating existing sequencing data ^44^ to define PAV. Strikingly, we found that maize lines without the SV showed increased TBN and had more upright leaves, while lines with the SV showed the opposite trend, and these results were significant by Wilcoxon rank-sum test (p-values TBN= 0.0078, LA= 0.043) at α = 0.05. Also, based on transcriptome data from the NAM Consortium, founder lines with the SV present in the *ereb184* promoter had lower gene expression (Wilcoxon rank-sum test *p*=0.029) in shoot tissue (Fig. 7c). These results show that SV in the *ereb184* promoter contributes to regulation of these pleiotropic, agronomic traits.

## Discussion

In maize, mutants that affect the ligule also affect tassel branch initiation and genes expressed at initiating ligules are also expressed at the boundary of other lateral organs ^10,11^ such as tassel branches. In this study, we leveraged this well-characterized maize genetics system to investigate the molecular underpinnings of pleiotropy: network rewiring around pleiotropic factors in two developmental contexts, the redeployment of TFs for analogous but different developmental processes, and *cis*-regulatory variation modulating pleiotropic loci. By strategically integrating context-specific biological data and multivariate GWAS models that exploit maximal diversity in TBN and LA, we identified new regulatory factors contributing to architectural pleiotropy in maize. Our approach can be applied in any genetic system to disassociate pleiotropic phenotypes through manipulation of *cis*-regulatory components and gene network connections.

Pleiotropy in crop phenotypes can limit productivity ceilings. For example, selection on a certain desired phenotype may come with unintended deleterious manifestations of another. This can happen when a gene controlling a target trait also functions in another developmental or physiological context. In recent years, the dramatic production gains of the 20th century have plateaued in the world’s most important cereal crops ^45^. Looking forward, step changes in crop improvement and sustainability will rely on targeted manipulation of regulatory pathways to fine-tune agronomic traits for enhanced plant productivity and resilience in dynamic environments ^46,47^. Central to this is the ability to predict and design highly-specific genetic changes at pleiotropic loci with minimal perturbation to the complex networks in which they reside. Therefore, knowledge of context-specific gene networks and functional *cis*-elements will enable greater precision in engineering or breeding optimal plant ideotypes.

In comparing predicted gene regulatory interactions between tassel branch and ligule development, both conservation and rewiring of the GRNs were observed. For example, KN1 and ZHD15 were maintained as hubs with conserved regulatory connections in both developmental contexts. Conservation around KN1 was not unexpected given its role as a master regulator of meristem maintenance. Identifying ZHD15 as a conserved hub was novel, but consistent with several lines of evidence in this study that implicate ZHDs as central players in architectural pleiotropy between tassel branching and leaf architecture. Alternatively, there was extensive rewiring of predicted network connections around LG2, which strongly controls TBN and LA in a pleiotropic manner. This suggests that re-deployment of LG2 in different developmental contexts was likely accompanied by loss and gain of regulatory connections to gene targets and targeting TFs. Perhaps LG2 serves a common function in setting up boundaries and its interactions with context-specific sub-networks shape the specific developmental programs.

We showed that these biological context-specific networks could be used to subset markers for multi-trait GWAS and identify new loci that contribute to architectural pleiotropy in TBN and LA. Using two different network-guided approaches we identified several gene candidates in and around significant SNP-trait associations. Strikingly, both approaches identified significant SNPs within two of the same TF-encoding genes, *zhd21* and *ereb184.* Both TFs are members of large families that are largely uncharacterized in maize and our results showed that *zhd* and *ereb* TFs were highly interconnected within the regulatory networks controlling tassel and leaf architecture. Furthermore, stacking *zhd* alleles with no observable phenotypes in these traits resulted in pleiotropic phenotypic expressions. This suggests functional redundancy among *zhd* genes and that further analyses of genetic interactions among *zhd* mutant alleles should provide insight into how to precisely manipulate plant architecture.

The vast stretch of intergenic space (∼100 kb) upstream of *ereb184* is sprinkled with regulatory signatures, including tissue-specific accessible chromatin regions, which potentially regulate its context-specific expression. The SV in its upstream promoter region, which is present in approximately half of the NAM founders, appears to control expression of *ereb184*; expression levels are significantly different in NAM lines with the SV compared to those that don’t. Presence or absence of the SV may contribute to differences in spatiotemporal expression patterns of *ereb184* and underlie the shifts in pleiotropy between TBN and LA. The origin of this SV is unclear. A blast search of the SV DNA sequence against a curated transposable element database revealed the presence of a probable LTR Gypsy retrotransposon (86% alignment identity) in the 3-prime region of the SV, as well as interspersed Helitron fragments. Differential chromatin accessibility between tassel primordia and shoot apex was evident in a regulatory region immediately proximal to the SV. Perhaps the SV is disrupting TF-binding interactions in the *ereb184* promoter or even interfering with long-range regulation through chromatin looping as suggested by a proximal Hi-C interaction.

Small effect loci for target traits are difficult to pinpoint with GWAS given that large trait effects generally dominate, which is compounded by the threshold for significance rising with marker density due to multiple correction. Targeted GWAS with a limited, but biologically informed marker set, has been demonstrated using pathway enrichment analysis ^48^ and chromatin accessible regions ^49^. Application of these methods to identify informative smaller SNP sets can enhance genomic predictions. Here, we showed that even SNP subsets identified with small numbers of genes prioritized through network analyses not only produced heritabilities similar to a genome-wide set of markers, but also enabled detection of small effect loci that were validated to contribute to both tassel and leaf architecture. Notably, we further showed that the context-specific networks from maize could be used to inform GWAS for LA in closely related sorghum. The analysis in sorghum identified *ereb114*, a paralog of *ereb184*, which was connected to it and *zhd21* in the network.

Our analyses support the utility of biologically informed, context-specific networks for guiding GWAS in various applications and for identifying loci that connect phenotypic traits. Results highlight various mechanisms by which expression of pleiotropic trait phenotypes are modulated, including through network interconnectedness of functionally redundant TF family members or SV in *cis*-regulatory components. We anticipate that this approach can be used widely to identify pleiotropic loci for manipulation in crop improvement.

## Supplementary Figure Legends

**Supplementary Fig. 1: Overview of biological samples collected for this study**

Representative images of hand-dissected tassel primordia from B73 control plants at **A**. stage 1, which captures branch meristem initiation and **B**. stage 2, which captures branch outgrowth. Scale bars = 1,000 μm. **C**. Longitudinal section through the shoot apical meristem (SAM) of a B73 control plant depicting the microscopic view of the shoot apex to the right. *In situ* hybridization with a *liguleless1* (*lg1*) anti-sense probe marks developing ligules. Two sections from the shoot apex were sampled: shoot apex 1 includes the SAM and cells pre-patterned to be ligule and shoot apex 2 excludes the SAM and enriches for developed ligule. Leaf primordia are designated according to plastochron number (P). White arrow denotes the pre-ligular band while black arrows indicate the developing ligule on P7 and successive leaf primordia.

**Supplementary Fig. 2: Principal component analysis of the expression dataset**

The image shows the three principal components in 3-dimensional space of the entire transcriptional dataset (*n*=140) generated from the two tassel primordia developmental stages and shoot apex sections. Genotypes are marked by different colors and tissue type by different symbols. The principal component analysis was calculated based on the top 500 dynamically expressed protein-coding genes.

**Supplementary Fig. 3: Dynamically expressed genes during tassel branching and ligule differentiation**

**A.** Volcano plot showing variation in gene expression associated with tassel branch development based on comparing between tassel stages 1 and 2 in B73 normal plants. Genes expressed higher in stage 2 are in blue and those in yellow are decreasing in expression. **B.** Volcano plot representing variation in gene expression associated with ligule differentiation based on the comparison between the shoot apex 1 and 2 samples in B73 normal plants. Genes expressed higher in shoot apex 2 are in blue and those in yellow are decreasing in expression. Only genes with FDR < 0.05 are plotted along the x- (log2 fold change) and y- (-log10 FDR) axis. Some classical maize genes are annotated.

**Supplementary Fig. 4: Many genes are differentially expressed in two developmental contexts during tassel branch outgrowth and ligule differentiation**

The heatmap displays relative gene expression profiles of annotated TFs and classical maize genes across both stages of tassel primordia and shoot apices sampled (n = 86). Genes were differentially expressed in the comparisons between tassel stages 1 and 2 and between shoot apex 1 and 2 with FDR < 0.05 (gray area of the Venn diagram). For each developmental context, DE genes that were up-regulated across developmental time are annotated with a blue box and down-regulated genes annotated with a yellow box.

**Supplementary Fig. 5: Maize developmental marker genes correspond to meristem identity and determinacy shifts across the tassel developmental gradient**

Expression trajectories of known maize developmental marker genes for meristem identity and determinacy are visualized to assess robustness of the five stage B73 normal tassel developmental gradient. **A**. Heatmap shows gene expression of selected maize inflorescence markers across the five tassel primordia stages in B73. **B.** Peak expression of classical maize genes associated with each stage of meristem development in B73 is shown on the left, and expression profiles for these marker genes in *liguleless2* and *ramosa1* mutant backgrounds are shown on the right. The *branched silkless1* profile is not shown in the mutants since it comes on after stage 2.

**Supplementary Fig. 6: Differences in gene expression of BR mutants compared to B73 controls in stage 2 tassels**

The ternary plot shows the relative expression of genes mis-expressed in common between the two BR mutants compared to B73 at stage 2. Each dot represents a gene and its coordinates indicate relative expression in the three genotypes.

**Supplementary Fig 7: Eigengene trajectories of co-expression modules from ‘tassel’ and ‘leaf’ networks**

The expression profile that best fits an average of all genes within a co-expression module is depicted as a Module Eigengene (ME). For each co-expression module in the **A.** ‘tassel’ and **B.** ‘leaf’ GCNs, MEs are depicted by dots in each sample. Each module showed a distinct expression profile, with certain modules characterized by peak expression at specific developmental stages and/or in certain mutant backgrounds.

**Supplementary Fig. 8: Density plots representing empirical distributions in comparison to whole genome *h^2^*and TBN and LA *h^2^***

**A.** TBN *h^2^*. Above 0 density represents the null distribution (purple arrow shows the 95th percentile) in comparison to *h^2^*explained by the selected co-expressed network modules (orange arrow) and whole genome *h^2^* (green arrow). Below 0 density represents null distribution (purple arrow shows the 95th percentile) in comparison to *h^2^*explained by the combined network motif (orange arrow) and whole genome *h^2^* (green arrow). B. LA *h^2^*. Upper panel represents the null distribution (purple arrow shows the 95th percentile) in comparison to *h^2^* explained by the selected co-expressed network modules (orange arrow) and whole genome *h^2^*(green arrow). Lower panel represents null distribution (purple arrow shows the 95th percentile) in comparison to *h^2^* explained by the combined network motif (orange arrow) and whole genome *h^2^* (green arrow)

**Supplementary Fig. 9: Depiction of statistical genetic interactions predicted for ereb184 with the top recurrent TFs within three-node network motifs**

Based on makers within the genomic windows defined as 土 2 kb from the transcriptional start site (TSS) and transcriptional termination site (TTS) of *ereb184 and the recurrent TFs from 3-node motifs*, statistical genetic interactions were predicted. These interactions are represented as links between genes located on a circular visualization of the maize genome. Grey and tan links represent significant interactions for the traits TBN, LA and PhPC2. Tan links highlight statistical interactions that are supported by the GRN data.

## Supplementary Tables

**Supplementary Table 1:** Gene expression in TPM across 140 RNA-seq samples.

**Supplementary Table 2:** DE genes with FDR < 5% derived from the comparison of B73 stage 2 tassel primordia vs. stage 1.

**Supplementary Table 3:** DE genes with FDR < 5% derived from the comparison of B73 shoot apex 2 vs. shoot apex 1.

**Supplementary Table 4:** GO terms associated with DE genes derived from the comparison of B73 stage 2 tassel primordia vs. stage 1.

**Supplementary Table 5:** GO terms associated with the DE genes derived from the comparison of B73 shoot apex 2 vs. shoot apex 1.

**Supplementary Table 6:** Genes that are DE between tassel stage 1 and stage 2 and shared between the BR mutants compared to B73 control.

**Supplementary Table 7:** Relative expression of genes commonly DE in the two BR mutants along the tassel stage 1 to stage 2 developmental gradient compared to the B73 control.

**Supplementary Table 8:** GO and TF enrichment analysis of genes commonly DE in the two BR mutants between tassel stage 1 and stage 2 development compared to the B73 control.

**Supplementary Table 9:** Summary of gene co-expression modules from the ‘tassel’ and ‘leaf’ context-specific networks.

**Supplementary Table 10:** Maize genes grouped based on the ‘tassel’ and ‘leaf’ gene co-expression modules.

**Supplementary Table 11:** Gene co-expression module preservation matrix between ‘tassel’ and ‘leaf’ networks.

**Supplementary Table 12:** GO terms overrepresented within ‘tassel’ and ‘leaf’ GCN modules.

**Supplementary Table 13:** GO terms overrepresented within ‘tassel’ and ‘leaf’ GCN preserved sub-modules.

**Supplementary Table 14:** TF enrichment analysis of the predicted regulators of *liguleless 2* in the leaf GRN shown in Figure 2c.

**Supplementary Table 15**: i) Summary statistics of the three-node network motifs annotated in the GRNs. ii) Top one-hundred recurrent TFs within the three-node network motifs.

**Supplementary Table 16:** GEBVs and BLUPs of the elected Ames diversity lines.

**Supplementary Table 17:** Pearson correlations of phenotypic traits: Growing degree days to silking (GDD_DTS), Growing degree days to anthesis (GDD_DTA), Predicted GEBV of Tassel Branch Number (TBN predicted), Predicted GEBV of Leaf Angle (LA predicted); Predicted GEBV of the three principal components of LA, TBN, and ERN (PhPC1 predicted, PhPC2 predicted, PhPC3 predicted),TBN, LA, PhPC1, PhPC2.

**Supplementary Table 18:** i) Module-trait correlation of the sixteen modules identified in the comprehensive GCN. ii) maize genes grouped based on the comprehensive GCN modules.

**Supplementary Table 19:** Gene co-expression module heritabilities (h2) for leaf angle (LA) and tassel branch number (TBN).

**Supplementary Table 20:** GWAS SNP-trait associations for LA, TBN, PhPC1 and PhPC2 from the single-trait approach using two sets of markers.

**Supplementary Table 21:** GO terms associated with genes proximal to significant SNP-trait associations from the single-trait GWAS for PhPC2.

**Supplementary Table 22:** Multi-trait GWAS SNP-trait associations for LA, TBN using two sets of markers.

**Supplementary Table 23:** Predicted direct gene targets of EREB184 and ZHD21 in the ‘leaf’ GRN.

**Supplementary Table 24:** Statistical genetic interaction predictions between *ereb184 and* top recurrent TFs within three-node network motifs.

**Supplementary Table 25:** Single-trait GWAS results for LA based markers subset around sorghum orthologs of the highly connected three-node motifs in the maize networks.

**Supplementary Table 26:** GO terms associated with genes in position Z of *ereb184*-controlled FFLs in the ‘leaf’ network.

## Data accessibility

Raw and processed data are available through NCBI Gene Expression Omnibus (GEO) database using the identification number GSE180593.

The manuscript utilizes data from various public datasets, including: conserved non-coding sequences ^44^ and unmethylated regions ^43^, which were downloaded from https://www.maizegdb.org, tassel inflorescence ATAC-seq and tassel chromatin interaction (Hi-C) (SRA ID: SRR10873334, SRR10873333, SRR10873339, SRR10873340, SRR10873341, SRR10873342, SRR10873343) ^41^, leaf chromatin interaction data (Hi-C) (SRA ID: SRR7889833, SRR7889834) ^42^.

## Acknowledgements

The authors would like to thank members of the Eveland, Lipka and Hake labs who assisted in tissue dissections and/or field phenotyping that contributed to this manuscript, namely Dr. Annis Richardson, Dr. Rajiv Parvathaneni, Dr. Samuel Leiboff, Dr. Matthew Murphy, Dr. Indrajit Kumar, Dr. Samuel Bonfim Fernandes, Ms. Zhonghui Wang, Ms. Adrianna Chepote, Dr. Yuguo Xiao, and Mr. Jake Sinkowitz. Also, a dear thanks to Dr. Todd Mockler and members of his team, Mr. Robert Lowery, Mr. Darren O’Brian, and Mr. Phil Ozersky, for their assistance in field phenotyping. We also thank Dr. Baoxing Song and Dr. M. Cinta Romay for providing the AGPv4 genome alignment BAM files for the resequencing data of the Maize 282 Association Panel. A special thank you to Kevin Reilly and his team in the Danforth Center Integrated Plant Growth Facility for their help with plant care as well as Noah Fahlgren and his team in the Danforth Data Science Core Facility with maintaining computational infrastructure used for this project. This work was funded by a National Science Foundation Plant Genome Research Project award # IOS-1733606 to ALE, AEL and SH.

## Authors’ contributions

ALE, AEL and SH designed the research. EB performed network analyses, data integration and interpretation. BRR performed genotype selection and multi-trait GWAS. MB and JY contributed to experimental design and performed molecular and genetics experiments. EB and JS performed genetic analyses of *zhd* alleles, EB and ALE wrote the paper with input from all authors. All authors contributed to editing the final manuscript, read and approved the final manuscript.

## Notes

### Competing Interest Statement

The authors have declared no competing interest.

### Summary of Updates

Manuscript condensed and revised. Results form figure six confirmed and updated.

