## Supplementary material for "Regulatory variation controlling architectural pleiotropy in maize": Methods

#### Plant material for RNA-seq experiments

Mutants were introgressed at least five times into the maize B73 inbred. All were grown along with B73 controls in environmentally-controlled growth chambers at the Danforth Center Integrated Plant Growth Facility with 14 hr days, 28° C / 24° C, 50% humidity and 450 μMol light. Plants were sown in cone trays (5 cm diameter, 11.5 cm depth, 142 ml total volume) in a Metromix 360-turface blend. At 14 Days After Sowing (DAS), seedlings were transplanted to larger pots (27 cm diameter, 24 cm depth, 14 L total volume; three plants per pot) with Berger 35% soil and 10 g of corn top dressing. In a trial experiment prior to tissue sampling, development of tassel primordia was tested and staged for uniformity across the different genotypes. Plants were staggered across a two week time period for tissue collections in identical conditions.

#### Tissue sampling and RNA extraction

Tassel primordia were hand-dissected and flash-frozen in liquid N. For each genotype, fifteen primordia were pooled per replicate for stage 1 and ten for stage 2 (Supplementary Fig. 1). Shoot apex 1 and 2 samples were collected from plants at 21 DAS. To collect shoot apex 2, whole plants were removed from the soil and whorls were removed until the developing ligule was ~.75cm from its node. Progressively younger leaf whorls (3-4 altogether) were removed at the connection to their node and a 2 mm section of leaf surrounding the ligule region was cut with a razor blade. From the remaining tissue, shoot apex 1 was collected by cutting an additional 2 mm section to include the base of the remaining developing leaves and the region including the shoot apical meristem. 2-3 individuals were collected per replicate and material was flash frozen immediately after dissection.

For each tissue type and developmental stage, four biological replicates were collected. Tissue samples were ground using a bead shaker with liquid nitrogen in a 2 ml tube with a 5 mm ceramic bead. RNA isolation from tassel material was performed using the PicoPure RNA isolation kit (ThermoFisher Scientific) according to manufacturer's instructions with the following adjustments: 40 μl and 60 μl of RNA extraction buffer were added respectively to stage 1 and stage 2 ground tassels. After 30 min incubation at 42°C, samples were centrifuged at 800 g for 2 min. Equal volume of 70% EtOH was added to samples and processed according to kit directions. On-column DNaseI treatment was performed per instructions using RNase-Free DNaseI kit (Qiagen) to remove residual DNA.

RNA isolation from shoot apex samples was performed using the Zymo quick-RNA plant kit according to manufacturer instructions with the following adjustments: Lysis buffer was added directly to the ground tissue and centrifuged, supernatant was added directly to the filtration column. DNaseI treatment was performed using the supplied DNaseI enzyme according to manufacturer instructions. All RNA quality was quantified using the NanoDrop One Spectrophotometer (Thermo Fisher Scientific) and using the RNA-6000 Pico chip from (Agilent) to ensure RNA integrity.

**RNA-seq libraries, sequencing and data analysis**

Poly(A)$^{+}$ RNA-seq library preparation and sequencing were outsourced to Novogene (USA). Libraries were multiplexed 12 per lane and sequenced using the Illumina HiSeq4000 platform with a 150-bp paired-end design. On average we generated more than 60 million paired-end reads per sample with quality score (Phred-score) > 30. Raw reads were processed to filter out low quality reads, adaptors or barcode remnants using *Cutadapt* v2.3 [1] and the wrapper tool *TrimGalore* v0.6.2 (http://www.bioinformatics.babraham.ac.uk/projects/trim_galore/) with default parameters except for --length 70 --trim-n --illumina. Clean reads were used to quantify the maize B73 AGPv4 gene models with *Salmon* v1.4.0 using the selective alignment method with a decoy-aware transcriptome [2]. A Salmon index was created using default parameters from the cdna fasta file (Zea_mays.AGPv4.cdna.all.fa) together with the genome reference fasta file (Zea_mays.AGPv4.dna.toplevel.fa) to generate the decoys for the selective alignment method. Maize reference files were downloaded from Ensembl Plants release 34 (ftp://ftp.ensemblgenomes.org/pub/plants/release-34/fasta/zea_mays/). Briefly, clean reads were mapped to the reference transcriptome using *Salmon quant* command with default parameters except for the options -l A --numBootstraps 100 --validateMappings. Expression levels were imported in R using the Bioconductor package *tximport* [3], summarized to gene level using the function *summarizeToGene()* and presented in TPM (Transcript Per kilobase Million). Overall gene expression levels between replicated samples (n=4) were highly related with correlation coefficients $r \geq 0.92$.

Differential expression analysis was performed using the Bioconductor package DESeq2 [4]. Pairwise contrasts were applied to compare mutant genotypes against equivalent normal samples. To test differences along the tassel developmental gradient attributable to a given genotype in comparison with B73, we set up an interaction design formula: ~ Genotype + Tissue + Genotype:Tissue. Genes were considered differentially expressed based on false discovery rate ≤ 0.05.

To standardize the relative expression of each gene across the three genotypes (*bin2*, *bri1*, B73 control), we normalized the expression values for each gene within the triad as follows: relative expression $bin2_{gene(i)}$ = TPM ($bin2_{gene(i)}$) / TPM ($bin2_{gene(i)}$) + TPM ($bri1_{gene(i)}$) + TPM ($\mathrm{B73}_{gene(i)}$); relative expression $bri12_{gene(i)}$ = TPM ($bri1_{gene(i)i}$) / TPM ($bin2_{gene(i)i}$) + TPM ($bri1_{gene(i)i}$) + TPM ($\mathrm{B73}_{gene(i)i}$); relative expression $\mathrm{B73}_{gene(i)}$ = TPM ($\mathrm{B73}_{gene(i)}$) / TPM ($bin2_{gene(i)}$) + TPM ($bri1_{gene(i)}$) + TPM ($\mathrm{B73}_{gene(i)}$).

Expression time (ET) was calculated using a smooth spline regression model [5] with the R function *bs()*. We fitted a b-spline (3-knot with three degrees of freedom) modeled on the first and second PC of the 500 most dynamically expressed genes across normal tassel development. Data points from mutant backgrounds were classified based on their location on the spline in relation to this model.

**Gene network analyses**

GCNs were built using the R package *WGCNA* (v.1.68) [6]. Expression data of protein coding genes were imported into R with the function *DESeqDataSetFromTximport()*. For each GCN we selected expressed genes based on row mean > 5 counts with the R function *rowMeans()* and normalized the count expression level of each gene according to the variance stabilizing transformation (VST) with the function *vst()* from *DESeq2* package. The Pearson correlation was used to select samples for the gene co-expression networks. Highly correlated biological replicates with $r \geq 0.92$ were retained and independently input in the network analyses. Based on the correlation coefficient, only one sample derived from the *lg1-R* stage 1 tassel (replicate 2) was excluded from the network analyses.

The soft power threshold was set to 6 for the ‘tassel’ and ‘leaf’ GCNs and to 7 for the combined network. Module detection was calculated via dynamic tree cutting using the function *blockwiseModules()* with the following parameters: type = signed, corType = bicor, minimum module size = 30, mergeCutHeight = 0.25. The parameter maxBlockSize for each network was set equal to the total number of expressed genes passing the mean cutoff as described above; 22,499 and 22,716, respectively, for ‘tassel’ and ‘leaf’ GCNs. The topographical overlap matrix (TOM) was calculated for each network using the function *TOMsimilarityFromExpr()* with parameters matching those used in the module detection. Networks were exported using the function *exportNetworkToCytoscape()* with parameters: weigh t= TRUE and threshold = 0.00. The R package *igraph* v1.2.4.1 [7] was used to build graphs from exported networks with the function *graph_from_data_frame()* and to calculate the graph statistics. Preserved modules between ‘tassel’ and ‘leaf’ GCNs were computed using the function *modulePreservation()* with

1,000 permutations. Gene sub-module preservation between networks was calculated using the R package *GeneOverlap* (v.1.28).

For the combined GCN generated from all samples, the module-to-sample association analysis was conducted evaluating the correlation between the module eigengene and samples of different developmental groups: i) tassel primordia at stage 1, ii) tassel primordia at stage 2, iii) shoot apex 1 and iv) shoot apex 2. In addition, we tested associations between module eigengene and three traits of interest: LA, TBN and TBA. A metafile was created where samples were categorized according to the four sample groups and three traits. The R function *cor()* and *corPvalueStudent()* were used to test the correlation between the module eigengene and the variables. Modules with $r > |0.8|$ were considered strongly correlated.

For the context-specific GRNs, a machine learning approach was applied to predict targets of known maize TFs using the Bioconductor package *GENIE3* [8]. Maize TFs were downloaded from the GRASSIUS repository (www.grassius.org) and overlapped with the expression matrices. TFs were set as “regulators” to infer their “targets” based on the gene expression abundance. *GENIE3* was run with parameters: treeMethod = "RF", nTrees = 1,000 and putative target genes were selected with a weight cutoff ≥ 0.005.

To determine enrichment of three-node subgraphs in the GRNs, we scanned for all possible three-node subgraphs and compared results with a set of randomized networks ($n$ = 1,000) with the same number of nodes and edges. This analysis was conducted with the R package *igraph* v1.2.4.1 using the following functions: *graph.full()* with $n$ equal to the number of genes in the context-specific GRNs, *triad.census()*, *cliques()* with min and max set to 3. Motif significance was determined by comparing the number of observed motifs with those found in the randomized networks. Genes involved in the fully connected three-node sub-graphs were selected and ranked based on their frequency.

**Transcription Factor and Gene Ontology enrichment analysis**

Maize TF and GO annotations were downloaded from GRASSIUS (www.grassius.org) [9] and GOMAP (doi.org/10.7946/P2M925) [10], respectively. The enrichment analysis was conducted with the Bioconductor package clusterProfiler [11] using the function *enricher()* with default parameters and cut-off of $q = 0.1$, where $q$ is the p-value adjusted for false discovery rate using the Benjamini-Hochberg correction. GO annotations were downloaded from the database QuickGO (https://www.ebi.ac.uk/QuickGO/).

### Germplasm selection and genotype data

A training set of 281 genotypes from the Goodman-Buckler diversity panel [12] was used to predict upper leaf angle (LA), tassel branch number (TBN), ear row number (ERN) and their corresponding principal components (PhPC1, PhPC2, PhPC3) in the Ames inbred diversity panel (a.k.a. the North Central Regional Plant Introduction Station (NCRPIS) panel) [13]. First, using publicly available multi-locations phenotypic data for the Goodman-Buckler panel [14,15], we fit a linear model with environment and genotype as fixed effects from which we obtained the best linear unbiased predictions (BLUPs) for each individual. The PCs of the three phenotype BLUPs, and all further phenotypic (Ph)PCs, were produced using the R function *prcomp()*. Data were centered and scaled before PhPC analysis. The genotype BLUPs of Goodman-Buckler panel phenotypes and PhPCs were used to train a genomic best linear unbiased prediction (GBLUP) [16] model that obtained predicted genomic estimated breeding values (GEBVs) for 2,534 Ames panel inbreds. Models for BLUPs, GBLUPs and GEBVs were conducted in R with the package *ASReml-R* [17]. Kinship matrices for GBLUP were produced as described in the 'Heritability' section below.

Genotypic data for the two diversity panels were downloaded from Panzea (www.panzea.org) and filtered for indels and non-biallelic markers. Missing data were imputed with the nearest neighbor method where distance is defined as linkage disequilibrium between two SNPs [18]. SNPs with low minor allele frequency were filtered using a 0.01 cut-off. Prior to analysis, genotypes used in this study were converted to AGPv4 coordinates using the tool CrossMap v0.3.7 [19] with the chain file from Gramene release 61 [20].

Nucleotide diversity, the average pairwise difference between all pairs of genotypes [21], was measured for each SNP using all genotypes for which GBS data were available. Marker filtering and nucleotide diversity calculations were achieved using *VCFtools* [22]

### Phenotypic data collections

Phenotypic data were collected at the University of Illinois, Urbana Champaign over three years (2018-2020). Each year, 425 Ames panel lines were randomly selected from across the distribution of the predicted PhPC1 values and planted. In addition, 75 lines from the Goodman-Buckler panel were planted (same lines each year) to ensure consistency across years. Lines were planted as single-row plots in mid-May each year; 28,000 seeds per acre with 30-inch row spacing. Field design was a randomized complete block design with two replicate blocks per year. Phenotypic observations were conducted during the first week of August after the majority of genotypes had flowered. Measurements of LA were taken from the leaf immediately above the uppermost ear. If no ear was present, we selected the fifth leaf below the flag leaf. Only plants with emerged tassels were phenotyped. The angle was measured from beneath the leaf from the horizontal axis, i.e., the stalk, to the midrib. An angle of 90° would indicate an entirely upright leaf, and an angle of 0° would be perpendicular to the stalk. TBN was conducted by counting every branch that originated from the tassel rachis. For each genotype three representative plants per plot were measured.

For each trait, we applied the mixed linear model to obtain BLUPs for each genotype:

$$Yijk = \mu + Gj + Ej + Bk(j) + \epsilon ijks \quad (3)$$

where, the phenotype $(Y)$ is explained by the $i^{\text{th}}$ genotype $(G)$ observed in the $k^{\text{th}}$ block $(B)$) nested in the $j^{\text{th}}$ year $(E)$. Individual plants within a plot are considered subsamples $(s)$.

After removing outliers, BLUPs for 1,064 and 1,072 genotypes for LA and TBN, respectively, were obtained.

**Heritability**

Prior to estimating heritability ( $\hat{h}^2$), SNP partitions were pruned with Plink 1.9 [23] to remove markers in LD of 0.7 or greater within a 50 bp window. The window was shifted five-bps and pruning repeated. For a given pruned SNP partition, a kinship matrix K was produced with the following model,

$$K = \frac{XX\prime}{n_p} (4)$$

where $X$ is the normalized SNP matrix, $X'$ is its transpose, and $n_p$ is the number or SNPs in the given partition. Narrow sense heritability $(\hat{h}^2)$ is estimated as $\frac{\widehat{\sigma_G}^2}{\widehat{\sigma_P}^2}$ where $\widehat{\sigma_G}^2$ is the additive genetic variance estimate and $\widehat{\sigma_P}^2$is the total phenotypic variance from model (2) when fitted using REML [24,25]. We generated null distributions by estimating $\hat{h}^2$ for 1,000 random gene sets using the SNPs found within their proximal regulatory region ($\pm$ 2 kb from TSS and TTS).

Random sets had an equal number of genes compared to the partition being tested. Genes in a given partition were removed from the entire genome-wide set before random selection. The software LDAK [26] was used to produce kinship matrices and estimate $\hat{h}^2$.

**Marker subsetting based on network analyses**

Genomic coordinates of gene sets derived from network analysis approaches (those from select co-expression modules and those most highly connected in three-node sub-graphs) were retrieved and imported in R. We used the Bioconductor package *GenomicRanges* [27] to select makers within the genomic windows defined as $\pm$ 2 kb from the TSS and the TTS of the co-expressed genes. Marker coordinates were intersected with the gene coordinates using the function *findOverlaps()* with the options type = within, ignore.strand = T.

**Genome-wide association studies**

We conducted single- and multi-trait association studies for LA and TBN. Single trait associations were conducted using Bayesian-information and Linkage-disequilibrium Iteratively Nested Keyway (BLINK) [28] conducted in GAPIT [29]. Before testing SNPs, the Bayesian information criteria (BIC) [30] was used to select the models with the optimal number of PCs using the Ames panel genome-wide pruned SNP dataset.

To test for pleiotropic associations, we utilized two approaches: i) BLINK, where the response variable was either the first or second PhPC of LA and TBN (PhPC1, PhPC2); and ii) multivariate extension of MLM (mvMLM), where the response was an *n*-by-*t* matrix with *n* being the number of observations and *t* the number of traits [31] conducted in GEMMA [32]. For mvMLM we conducted a leave-one-chromosome-out kinship approach [33]. Since kinship was chromosome specific the BIC optimal number of PCs was considered on a chromosome specific basis. Multiple testing correction was conducted based on the number of SNPs in a given partition using the Benjamini & Hochberg false discovery rate (FDR) procedure [34]. Genes with SNPs with an FDR-adjusted P-value below 0.2 were considered for further analysis.

**Statistical analysis of *ereb184* genetic interactions**

We tested for epistatic interactions by modifying the unified MLM as follows:

$$Y = Q\gamma + S_1\alpha_1 + S_2\alpha_2 + S_1S_2\beta + Z\mu + \varepsilon$$

where $\boldsymbol{Y}$ is *n* vector of phenotype BLUPs with *n* being the number of observations; $\boldsymbol{Q}$ is the *n-by-(p+1)* incidence matrix corresponding to the intercept, as well as *p* fixed effect covariates (i.e., principal components) accounting for subpopulation structure; $S_1$ is an *n-by-1* incidence vector for the peak associated SNP from *ereb184*; $S_2$ is an *n-by-1* incidence vector for the testing SNP and $S_1S_2$ their interaction; $\alpha_1$ is the additive effect of the peak associated SNP; $\alpha_2$ is the additive effect of the testing SNP; $\beta$ is the additive x additive epistatic effect between the peak associated SNP and the testing SNP; $\boldsymbol{Z}$ is an *n-by-n* incidence matrix relating u to $\boldsymbol{Y}$; $\mu$ ~ *N(0,*

$2K\sigma_G^2)$; and $\varepsilon \sim MVN(0, I\sigma_e^2)$ is the residual error with variance with *I* being the identity matrix and $\sigma_e^2$ the residual variance. The peak-associated SNP and the testing SNP are treated as fixed effects. The model was performed so that $S_2$ was a SNP assigned to a motif gene and run for each SNP in the motif gene partition including those assigned to *ereb184 that were not* $S_1$. The model was run in *ASReml-R* [17].

**Sorghum LA association study**

We retrieved sorghum orthologs based on [35], which identified 11,000 sorghum-maize syntenic orthologs (https://figshare.com/articles/dataset/Grass_Syntenic_Gene_List_sorghum_v3_maize_v3_4_with_teff_and_oropetium_v2/7926674/1). Of the 200 maize TFs within our top-ranked network motif connectedness, we identified 146 sorghum-maize syntenic orthologs. Sorghum BTx623 reference (version 3) gene coordinates were retrieved from the GFF Sbicolor_454_v3.1.1.gene.gff3 (Phytozome v12.1). Sorghum has larger LD blocks than maize [36] therefore for each sorghum-maize syntenic ortholog, we extended the gene coordinates of ±10 kb from TSS and TTS, respectively using the R package *GenomicRanges* and the function *start()* and *end()*. These sorghum orthologs with extended coordinates were used to subset proximal makers using the Bioconductor function *findOverlaps()* with the option type="within" and ignore.strand=T.

Sorghum LA phenotype data were previously collected for 296 individuals from the Sorghum Association Panel (SAP) [37] from the leaf below the flag leaf [38]. SAP GBS data [39] were filtered at MAF 0.05. All analyses in sorghum were conducted according to the methods described above for maize.

***ereb184* SV analysis in the Goodman-Buckler panel**

We retrieved existing sequencing data[40] from the Goodman-Buckler panel aligned to maize B73 AGPv4. Using Samtools v1.9, we extracted the total number of reads flagged as Q30 aligned to the genomic region defined as 1:286721317-286726162. To identify local differences within the defined region, we divided it into five equal-sized bins (969 bp) and recorded the number of aligned reads for each bin. The presence/absence of the structural variation (SV) was calculated as the ratio between the number of mapped reads in the bin and the total number of reads mapped to the entire region.

**Analysis of *zhd* UniformMu insertion lines**

The *zhd1* (Mu ID: *mu1022277*, AGPv4 coordinates: Chr4:12135894-12135902), *zhd21-1* (Mu ID: *mu1056071*, AGPv4 Chr3:137588503..137590511) and *zhd21-2* (Mu ID: *mu1018735*, AGPv4 coordinates: Chr3:137588828-137590836) alleles were isolated in the W22 inbred line carrying exonic Mutator (Mu) transposon insertions as part of the UniformMu transposon collection [41]. *Zhd* alleles were backcrossed into the W22 inbred line for at least two generations. Individual homozygous mutant alleles were grown at the Danforth Center Field Research Site during the 2023 season, utilizing a two-row design with a spacing of 2.5 feet between rows and 3 feet between ranges. Each genotype was represented by 40 plants. TBN and LA measurements were collected as described above.

Segregating populations of *zhd1* and *zhd21-2* mutant alleles were generated by genetic crosses. Maize plants were grown at North Carolina State Method Road Greenhouse under 16 hours of supplemental light/8 hours dark, and relative temperatures of 29.4°C day and 23.9°C night, and 12 inch pots in Metro-Mix 830-F3B (SunGro Horticulture). Genomic DNA was isolated from leaf tissue and gene-specific and transposon multiplex primer PCR was performed under standard conditions with 2X GoTaq Green Master Mix (Promega) with 5% DMSO (v/v). The insertions were confirmed, as was co-segregation of the phenotypes with the insertions, by PCR analysis using primers at the *zhd1* (CTCCTGGGGTTTGCAATTGC; GTGTGCATCATGTTCAGCGG) and *zhd21* (TTGTTGCAGCGTGAGACAGG; AGAAATCCATGGAGACTCCGC) loci in combination with Mu-TIR primer (AGAGAAGCCAACGCCAWCGCCTCYATTTCGTC). Tassel and leaf phenotypic data were collected from a population that segregated 1:1:1:1 for the following genotypes: *zhd1/+*; *zhd21-2/+*, *zhd1/zhd1*; *zhd21-2/+*, *zhd1/+*; *zhd21-2/zhd21-2*, *zhd1/zhd1*; *zhd21-2/zhd21-2*. Phenotypic data were not collected on double mutant plants due to delayed maturation. The blade/sheath boundary from mature leaves was scanned on each side with an Epson V600 flatbed scanner. Blade angle was quantified from scanned images of leaves with the angle tool and “measure” function in ImageJ. Tassel traits were scored by the following definitions: TBN, the number of basal long branches bearing only spikelet pairs.

**ATAC-seq libraries and data analyses**

Shoot apex 2 samples were collected as described above and flash-frozen. Tissue was ground in liquid N and ~0.2-0.3 g aliquoted into a 15 mL falcon tube. Ground tissue was resuspended in 4 mL of 1x nuclei isolation buffer (16 mM HEPES; pH8, 200 mM sucrose, 0.8 mM MgCl2, 4 mM KCl, 32 % Glycerol, 0.25% Triton X100, 1x complete protease inhibitor, 0.1% 2-ME, 0.1 mM

PMSF) very gently at 4°C for 20 minutes and then filtered through 2 sheets of mira cloth. The resulting elutent (~3 mL) was split equally into two 2 mL eppendorf tubes and centrifuged at 1,000 x g for 15 minutes. After discarding the supernatant, nuclei pellets in the two eppendorf tubes were resuspended in 400 µL of 1x tagmentation buffer, combined into a single tube and centrifuged at 1,000 x g for 5 minutes as a wash step (total 2 x washes). The nuclei were resuspended in 100 µL of 1x tagmentation buffer and observed under a microscope (2% acetocarmine stain) to check nuclear integrity. Nuclei were counted using a hemocytometer, and approximately 50,000 nuclei were used for tagmentation. Tagmentation was performed using the Illumina DNA Library Prep Kit (FC-121-1031) and Index Kit (FC-121-1011) with 2.5 µL Tn5 enzyme, 2.5 µL of 2x tagmentation buffer (20 mM Tris Base, 10 mM MgCl2, 20% v/v dimethylformamide), and 20 µL of nuclei for each sample and incubated for 1 hr at 37°C. To each reaction, added 22 µL of H2O, 2.5 µL 10% SDS, and 0.5 µL of Proteinase K and incubated at 55 °C for 1 hr. Tagmented libraries were purified using the Zymo clean and concentrator kit, eluting with resuspension buffer. DNA was quantified using Qubit. 25 ng of tagmented DNA was combined with 2.5 µL of both index primers, 7.5 µL PCR master mix, 2.5 µL NPM and filled to 25 µL with RSB buffer. Libraries were amplified with 11 cycles. Libraries were diluted to 50 µL and cleaned up with a two-sided Ampure XP bead size selection. A 0.5:1 bead:sample ratio followed by a 1.2:1 bead:sample ratio was used to select ~200-1000 bp libraries. Partial lane sequencing was performed using Illumina Novaseq 6000 platform with a paired-end 150 bp reads design. For each of the two biological replicates more than 89 million mapped paired-end reads were generated.

For data analysis, raw ATAC-seq reads were trimmed as described for the RNA-seq data with the additional parameters --stringency 1 -q 20. Cleaned reads were mapped to the maize AGPv4 genome using bowtie2 v2.4.5 [42] with default parameters except for --very-sensitive -X 2000 --dovetail. Reads mapped to mitochondria and chloroplast together with PCR duplicates and low mapping quality reads (mapping score < 10) were removed. Peaks were called using MACS2 v2.1.2 [43] with the parameters -f BAMPE --shift -100 --extsize 200 --nomodel -B --SPMR -g 2106338117. Consensus peaks were generated using the Bioconductor package *GenomicRanges* [27]. BigWig files were generated using bamCoverage with the parameters --binSize 1 --normalizeUsing RPKM.

**Methods References**

15. Tian, F. *et al.* Genome-wide association study of leaf architecture in the maize nested association mapping population. *Nat. Genet.* **43**, 159–162 (2011).
16. VanRaden, P. M. Efficient methods to compute genomic predictions. *J. Dairy Sci.* **91**, 4414–4423 (2008).
17. Butler, D. G., Cullis, B. R., Gilmour, A. R., Gogel, B. J. & Thompson, R. *ASReml-R Reference Manual Version 4*. (2017).
18. Money, D. *et al.* LinkImpute: Fast and Accurate Genotype Imputation for Nonmodel Organisms. *G3* **5**, 2383–2390 (2015).
19. Zhao, H. *et al.* CrossMap: a versatile tool for coordinate conversion between genome assemblies. *Bioinformatics* **30**, 1006–1007 (2014).
20. Jiao, Y. *et al.* Improved maize reference genome with single-molecule technologies. *Nature* (2017) doi:10.1038/nature22971.
21. Nei, M. & Li, W. H. Mathematical model for studying genetic variation in terms of restriction endonucleases. *Proc. Natl. Acad. Sci. U. S. A.* **76**, 5269–5273 (1979).
22. Danecek, P. *et al.* The variant call format and VCFtools. *Bioinformatics* **27**, 2156–2158 (2011).
23. Chang, C. C. *et al.* Second-generation PLINK: rising to the challenge of larger and richer datasets. *GigaScience* vol. 4 Preprint at https://doi.org/10.1186/s13742-015-0047-8 (2015).
24. Gilmour, A. R., Thompson, R. & Cullis, B. R. Average Information REML: An Efficient Algorithm for Variance Parameter Estimation in Linear Mixed Models. *Biometrics* **51**, 1440–1450 (1995).
25. Lee, S. H. & van der Werf, J. H. J. An efficient variance component approach implementing an average information REML suitable for combined LD and linkage mapping with a general complex pedigree. *Genet. Sel. Evol.* **38**, 25–43 (2006).
26. Speed, D. *et al.* Reevaluation of SNP heritability in complex human traits. *Nat. Genet.* **49**, 986–992 (2017).
