## Supplementary Information for "Regulatory variation controlling architectural pleiotropy in maize"

Table of Contents

Supplementary Note………………………………………………………………………….. 2

Supplementary Fig. 1: Overview of biological samples collected for this study…………… 5

Supplementary Fig. 2: Principal component analysis of the expression dataset………….. 6

Supplementary Fig. 3: Dynamically expressed genes during tassel branching and ligule differentiation……………………………………………………………………………… 7

Supplementary Fig. 4: Many genes are differentially expressed in two developmental contexts during tassel branch outgrowth and ligule differentiation………………………….. 8

**Supplementary Note**

***This supplementary note along with references describes the biology behind the mutants that we are using in this study.***

The *liguleless1 (lg1)* gene encodes a SQUAMOSA BINDING PROTEIN (SBP) TF and loss-of-function mutants completely abolish ligule and auricle formation, yet maintain a clear blade-sheath boundary [1,2]. In *liguleless2* (*lg2*) mutants, the blade-sheath boundary is more diffuse and reduced auricle tissue is retained at the leaf margins. This suggests that LG2, a bZIP TF, may be involved in defining the blade-sheath boundary [3] and that *lg1* is required for proper ligule differentiation [4].

During early tassel development, the indeterminate inflorescence meristem initiates a defined number of indeterminate axillary branch meristems at its flanks, which produce the long tassel branches, and then abruptly shifts to initiating short branches [5]. The *ramosa1* (*ra1*) gene, encoding a C2H2 TF specific to panicoid grasses, controls this process by imposing determinate fate on axillary meristems in a spatiotemporal manner [6]. The RA1 protein accumulates adaxial to the base of axillary meristems that will produce short branches [6]. Prior to RA1 expression, the LG1 protein accumulates in an overlapping domain at the base of the long branch meristems, but not the short ones, and RA1 can bind and negatively modulate *lg1* expression based on ChIP-seq and transcriptome studies [7].

Since RA1 is an inflorescence-specific TF, its regulation of *lg1* is not present in developing ligules. Similarly, WAVY AURICLE BLADE1 (WAB1), a TCP family TF, is a tissue-specific factor that promotes expression of *lg1* in the tassel. In *Wab1* gain-of-function mutants, LG1 over-accumulates at the base of tassel branches resulting in wider branch angles and is ectopically expressed in the leaf, causing aberrant ligule tissue in the sheath [8]. Loss-of-function *wab1* mutants (allelic to *branch angle defective1* (*bad1*)), have more upright tassel branches [8,9]. Another branching mutant, *ramosa2* (*ra2*), which encodes a LATERAL ORGAN BOUNDARY (LOB) domain TF [10], acts upstream of *wab1*/*bad1* and in a different pathway than *lg1* [9]. Tassel branches of *ra2* mutants are very upright.

RNAi knockdown lines of the BR receptor kinase, BRASSINOSTEROID INSENSITIVE 1 (BRI1), an initiator of BR signaling, and BR INSENSITIVE 2 (BIN2), a GSK3-like kinase and negative regulator of BR transcriptional response, show opposite architectural phenotypes. The *bri1-RNAi* mutant significantly knocked down *zmbri1* expression as well as expression of four other paralogs, causing decreased BR signaling; ligule and auricles are abnormal and leaves were more upright, and more upright tassel branches [11]. *bin2-RNAi* plants have wide open leaves due to expanded auricles and long, open tassel branches [12].

| **mutants used in network analyses** | **allele** |
|---|---|
| *brassinosteroid insensitive2 (bin2)* | *bin2*-RNAi |
| *brassinosteroid insensitive1* (*bri1*) | *bri1*-RNAi |
| *feminized, upright, narrow* (*fun*) | *fun1-1* |
| *liguleless1* (*lg1*) | *lg1-R* |
| *liguleless2* (*lg2*) | *lg2-R* |
| *ramosa1* (*ra1*) | *ra1-R* |
| *ramosa2* (*ra2*) | *ra2-R* |
| *wavy auricle blade1 (wab1)* | *wab-rev, Wab-1* (dominant) |

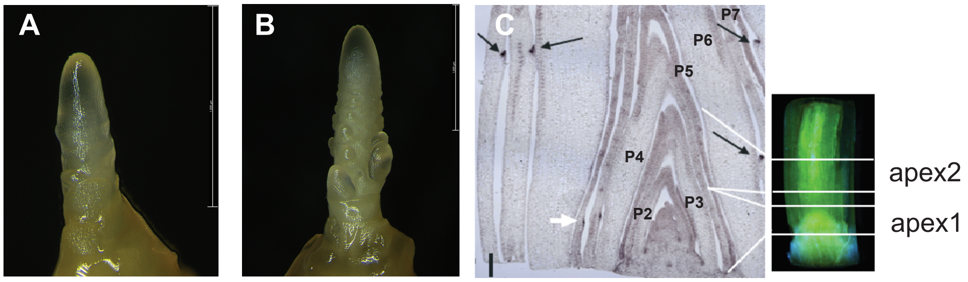

**Supplementary Fig. 1: Overview of biological samples collected for this study**
Representative images of hand-dissected tassel primordia from B73 control plants at **A.** stage 1, which captures branch meristem initiation and **B.** stage 2, which captures branch outgrowth. Scale bars = 1,000 µm. **C.** Longitudinal section through the shoot apical meristem (SAM) of a B73 control plant depicting the microscopic view of the shoot apex to the right. *In situ* hybridization with a *liguleless1* (*lg1*) anti-sense probe marks developing ligules. Two sections from the shoot apex were sampled: shoot apex 1 includes the SAM and cells pre-patterned to be ligule and shoot apex 2 excludes the SAM and enriches for developed ligule. Leaf primordia are designated according to plastochron number (P). White arrow denotes the pre-ligular band while black arrows indicate the developing ligule on P7 and successive leaf primordia.

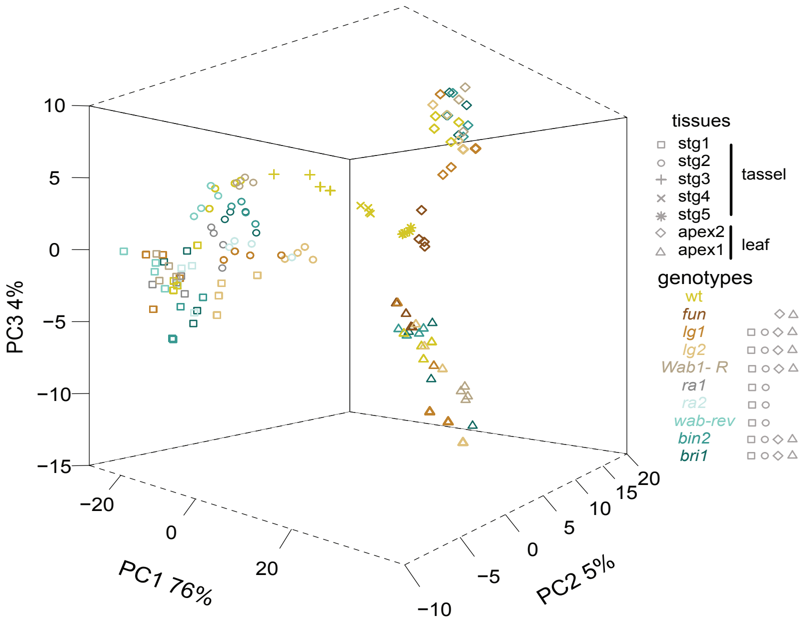

**Supplementary Fig. 2: Principal component analysis of the expression dataset**
The image shows the three principal components in 3-dimensional space of the entire transcriptional dataset (*n*=140) generated from the two tassel primordia developmental stages and shoot apex sections. Genotypes are marked by different colors and tissue type by different symbols. The principal component analysis was calculated based on the top 500 dynamically expressed protein-coding genes.

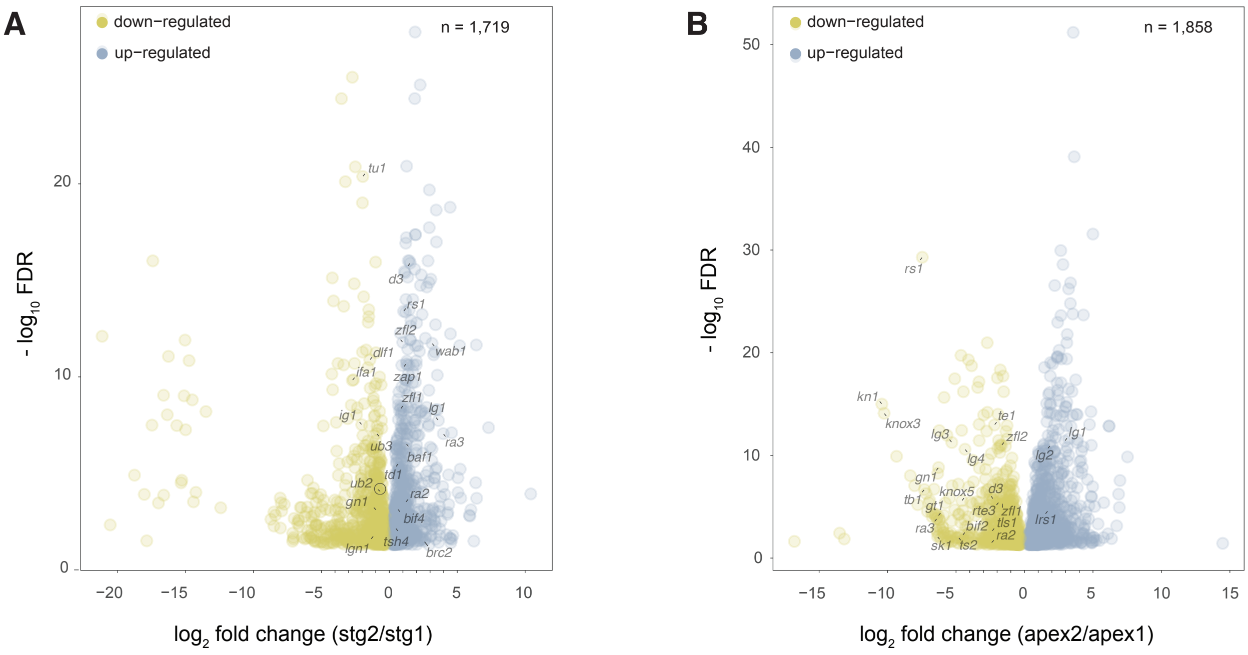

**Supplementary Fig. 3: Dynamically expressed genes during tassel branching and ligule differentiation**
**A**. Volcano plot showing variation in gene expression associated with tassel branch development based on comparing between tassel stages 1 and 2 in B73 normal plants. Genes expressed higher in stage 2 are in blue and those in yellow are decreasing in expression. **B**. Volcano plot representing variation in gene expression associated with ligule differentiation based on the comparison between the shoot apex 1 and 2 samples in B73 normal plants. Genes expressed higher in shoot apex 2 are in blue and those in yellow are decreasing in expression. Only genes with FDR < 0.05 are plotted along the x- ($log_2$ fold change) and y- ($-log_{10}$ FDR) axis. Some classical maize genes are annotated.

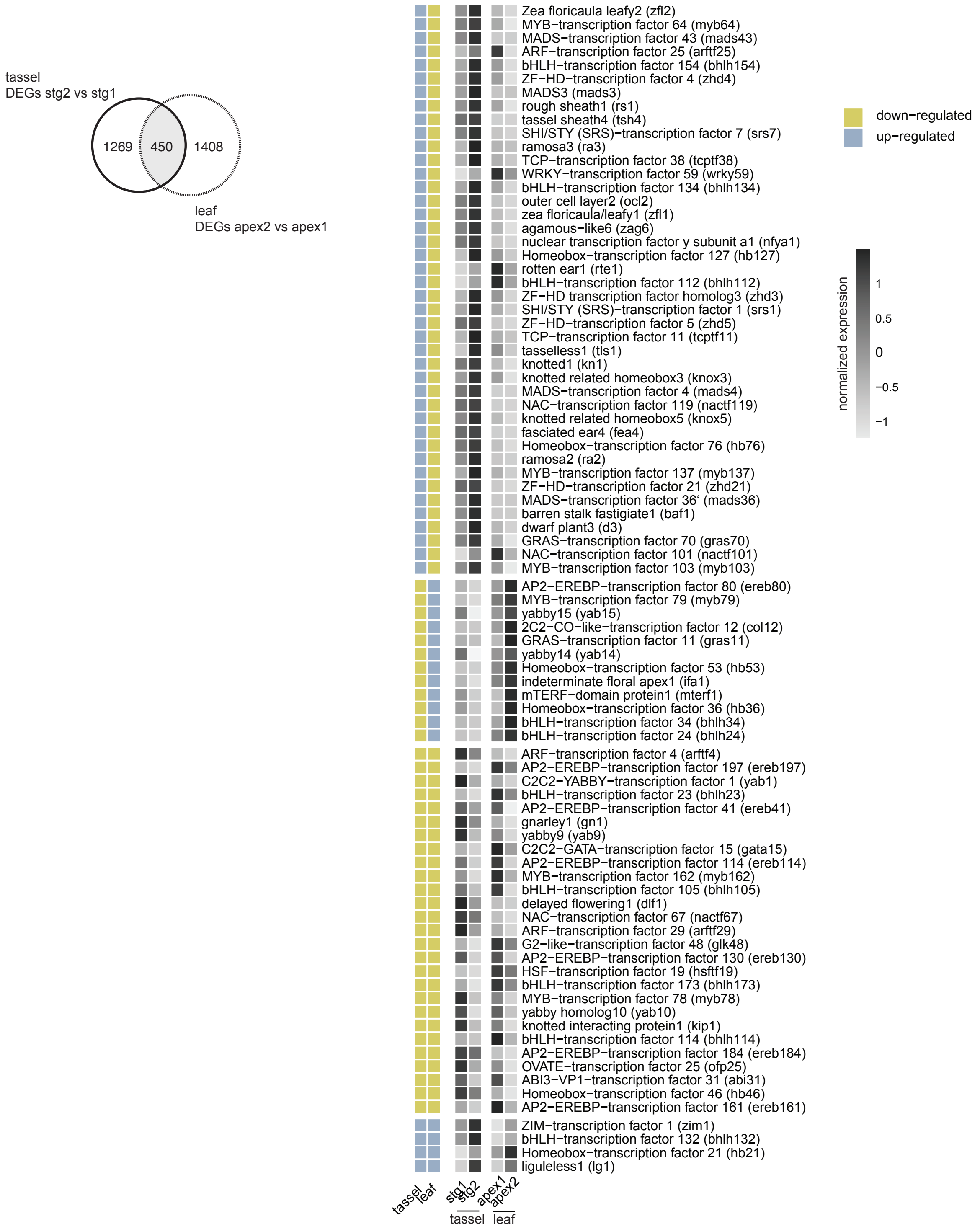

**Supplementary Fig. 4: Many genes are differentially expressed in two developmental contexts during tassel branch outgrowth and ligule differentiation**

The heatmap displays relative gene expression profiles of annotated TFs and classical maize genes across both stages of tassel primordia and shoot apices sampled (n = 86). Genes were differentially expressed in the comparisons between tassel stages 1 and 2 and between shoot apex 1 and 2 with FDR < 0.05 (gray area of the Venn diagram). For each developmental context, DE genes that were up-regulated across developmental time are annotated with a blue box and down-regulated genes annotated with a yellow box.

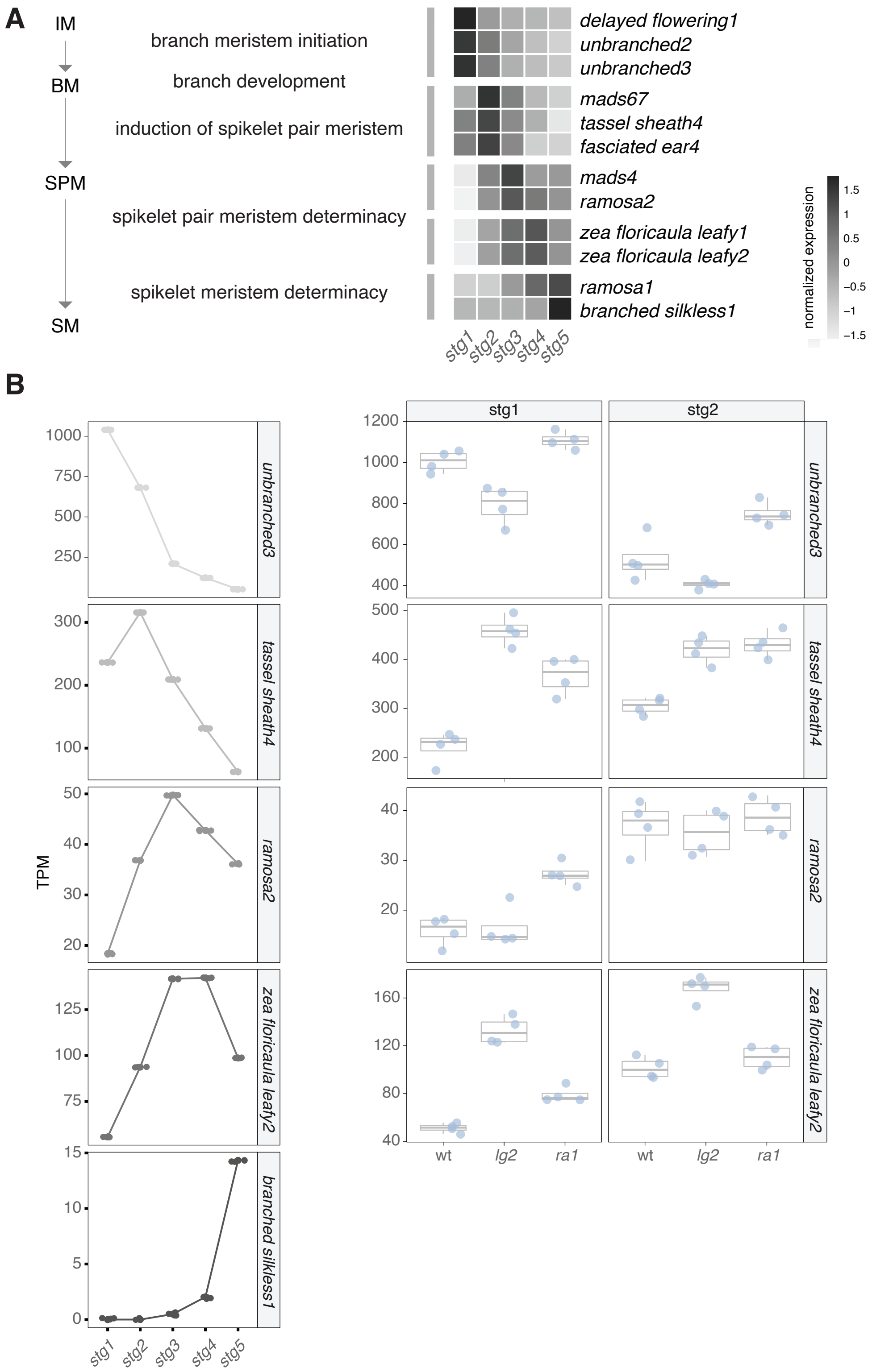

**Supplementary Fig. 5: Maize developmental marker genes correspond to meristem identity and determinacy shifts across the tassel developmental gradient**

Expression trajectories of known maize developmental marker genes for meristem identity and determinacy are visualized to assess robustness of the five stage B73 normal tassel developmental gradient. **A**. Heatmap shows gene expression of selected maize inflorescence markers across the five tassel primordia stages in B73. **B**. Peak expression of classical maize genes associated with each stage of meristem development in B73 is shown on the left, and expression profiles for these marker genes in *liguleless2* and *ramosa1* mutant backgrounds are shown on the right. The *branched silkless1* profile is not shown in the mutants since it comes on after stage 2.

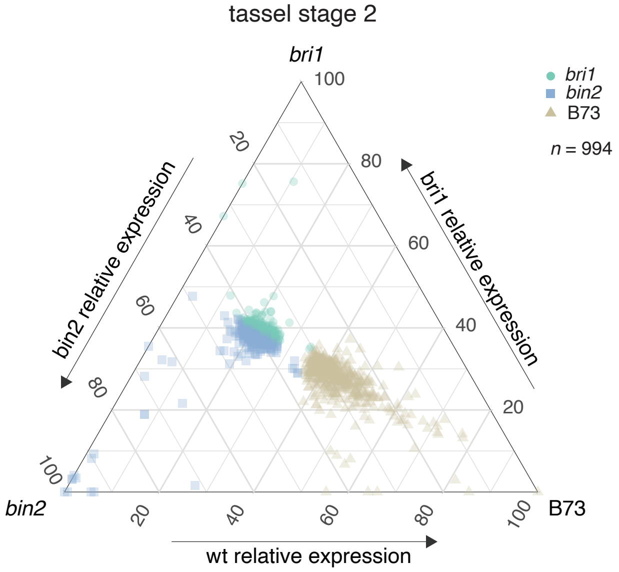

**Supplementary Fig. 6: Differences in gene expression of BR mutants compared to B73 controls in stage 2 tassels**

The ternary plot shows the relative expression of genes mis-expressed in common between the two BR mutants compared to B73 at stage 2. Each dot represents a gene and its coordinates indicate relative expression in the three genotypes.

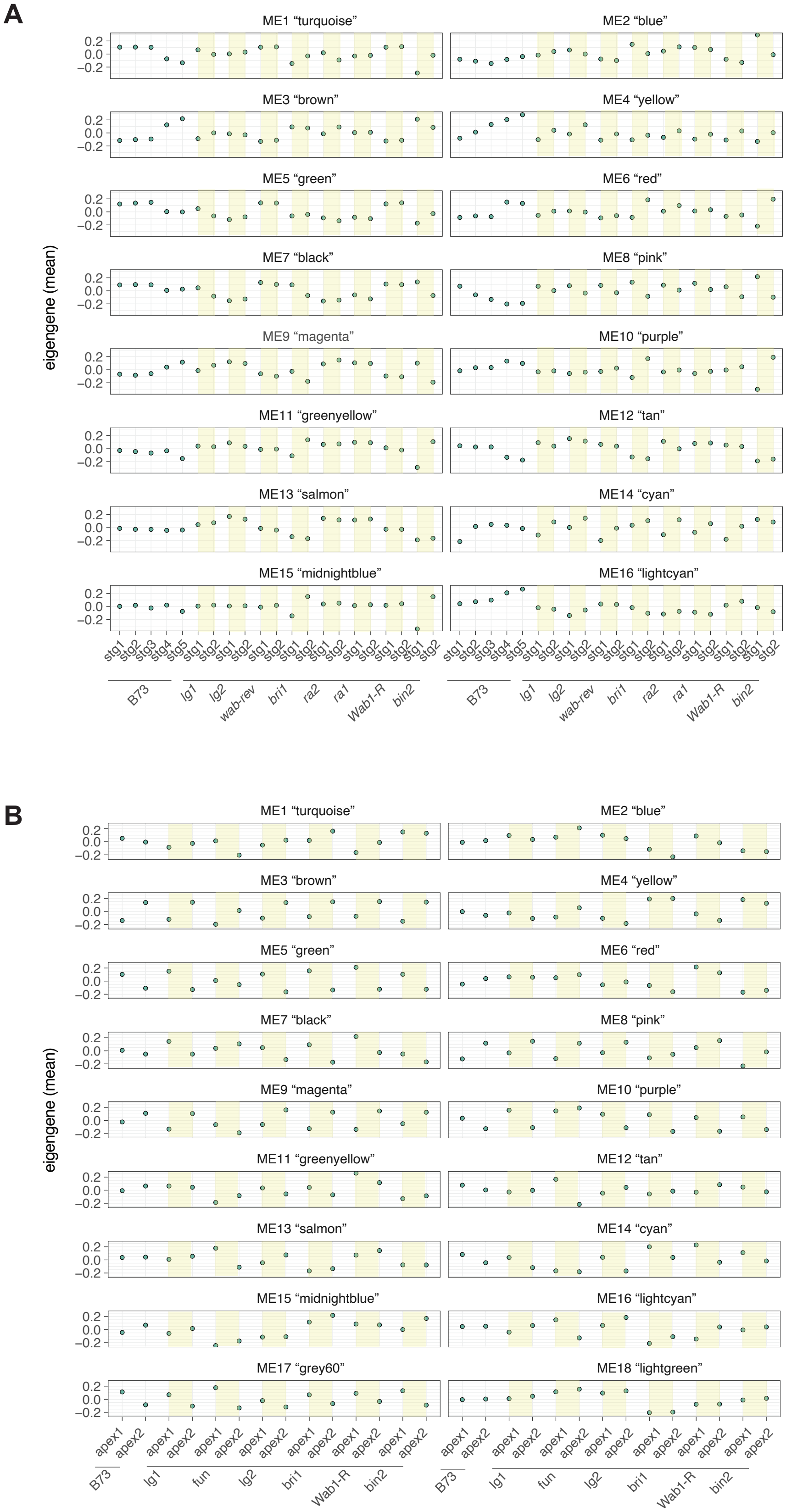

**Supplementary Fig 7: Eigengene trajectories of co-expression modules from ‘tassel’ and ‘leaf’ networks**
The expression profile that best fits an average of all genes within a co-expression module is depicted as a Module Eigengene (ME). For each co-expression module in the **A**. ‘tassel’ and **B**. ‘leaf’ GCNs, MEs are depicted by dots in each sample. Each module showed a distinct expression profile, with certain modules characterized by peak expression at specific developmental stages and/or in certain mutant backgrounds.

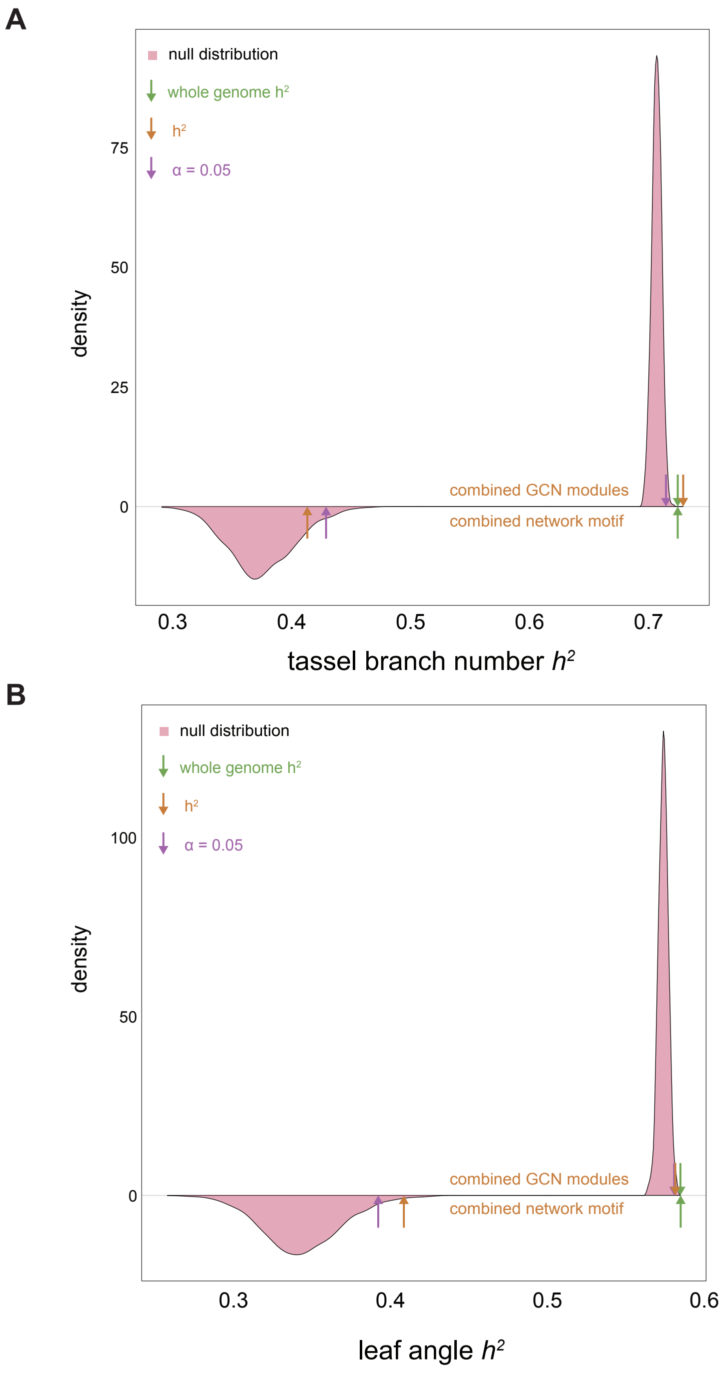

**Supplementary Fig. 8: Density plots representing empirical distributions in comparison to whole genome $h^2$ and TBN and LA $h^2$**

**A**. TBN $h^2$. Above 0 density represents the null distribution (purple arrow shows the 95$^{th}$ percentile) in comparison to $h^2$ explained by the selected co-expressed network modules (orange arrow) and whole genome $h^2$ (green arrow). Below 0 density represents null distribution (purple arrow shows the 95*th* percentile) in comparison to $h^2$ explained by the combined network motif (orange arrow) and whole genome $h^2$ (green arrow). **B**. LA $h^2$. Upper panel represents the null distribution (purple arrow shows the 95$^{th}$ percentile) in comparison to $h^2$ explained by the selected co-expressed network modules (orange arrow) and whole genome $h^2$ (green arrow). Lower panel represents null distribution (purple arrow shows the 95*th* percentile) in comparison to $h^2$ explained by the combined network motif (orange arrow) and whole genome $h^2$ (green arrow).

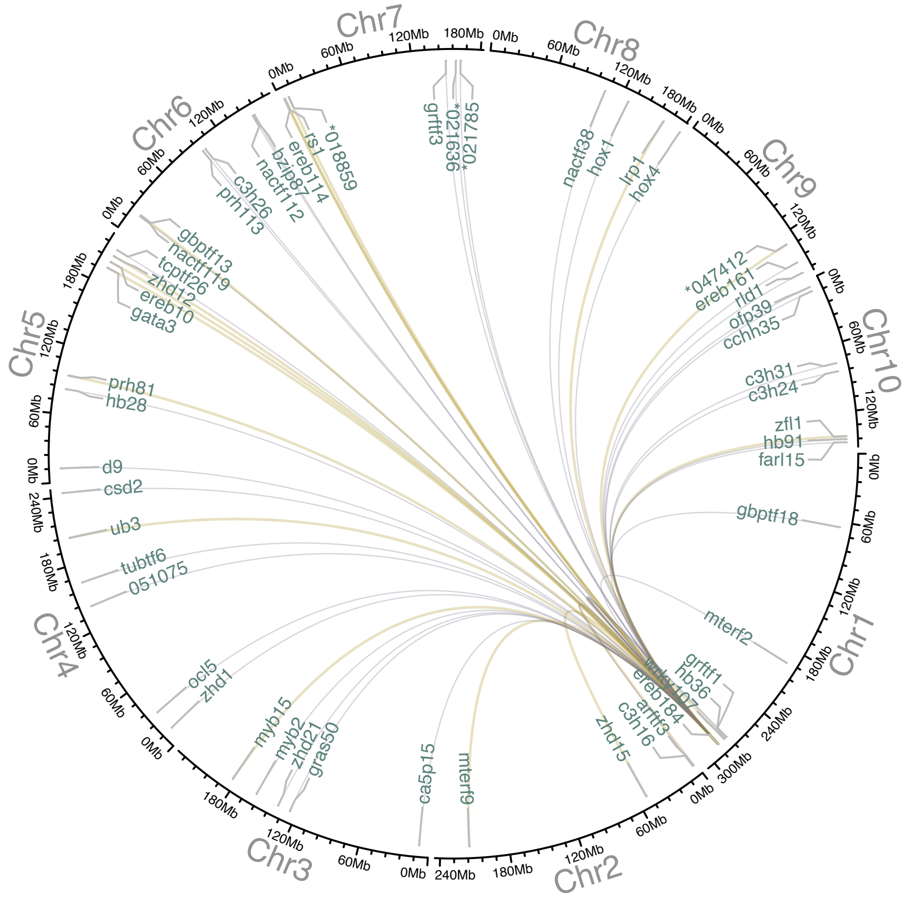

**Supplementary Fig. 9: Depiction of statistical genetic interactions predicted for *ereb184* with the top recurrent TFs within three-node network motifs**

Based on makers within the genomic windows defined as ± 2 kb from the transcriptional start site (TSS) and transcriptional termination site (TTS) of *ereb184* and the recurrent TFs from 3-node motifs, statistical genetic interactions were predicted. These interactions are represented as links between genes located on a circular visualization of the maize genome. Grey and tan links represent significant interactions for the traits TBN, LA and PhPC2. Tan links highlight statistical interactions that are supported by the GRN data.
